# Recent exposure to environmental stochasticity does not determine the resilience of natural populations

**DOI:** 10.1101/2022.04.28.489852

**Authors:** James Cant, Pol Capdevila, Maria Beger, Roberto Salguero-Gómez

**Affiliations:** Centre for Biological Diversity, University of St Andrews, St Andrews, KY16 9TH, United Kingdom; School of Biology, Faculty of Biological Sciences, University of Leeds, Woodhouse Lane, Leeds, LS2 9JT, United Kingdom; School of Biological Sciences, University of Bristol, 24 Tyndall Ave, BS8 1TQ, Bristol, UK; Department of Zoology, University of Oxford, 11a Mansfield Road, Oxford, OX1 3SZ, United Kingdom; Centre for Biodiversity and Conservation Science, School of Biological Sciences, University of Queensland, Brisbane, QLD 4072, Australia; Max Planck Institute for Demographic Research, Konrad Zuße Straße 1, 18057 Rostock, Germany

## Abstract

Escalating climatic and anthropogenic pressures expose ecosystems worldwide to increasingly frequent disturbances. Yet, our ability to forecast the responses of natural populations to these disturbances is impeded by a limited understanding for how exposure to stochastic environments shapes population resilience. Instead, the resilience, and vulnerability, of natural populations to ongoing global change is often presumed based on their contemporary exposure to environmental stochasticity. To test the validity of this assumption, we investigated the association between the resilience attributes (e.g., resistance and recovery) of natural animal and plant populations, and measures of local environmental stochasticity (e.g., spectral frequency and abiotic range); collating data from 2,242 populations across 369 animal, plant, and algal species. Unexpectedly, recent abiotic stochasticity regimes from the past 50 years do not predict the inherent ability of populations to resist or recover from disturbances. Instead, population resilience is strongly affected by phylogenetic relationships among species, with survival and developmental investments shaping their responses to stochastic regimes. Contrary to the classical assumption that exposure to recent environmental shifts confers a greater ability to cope with current and future global change, our findings suggest that population resilience is a consequence of evolutionary processes and/or deep-time environmental regimes.

**Significance statement:** Populations that currently endure more variable abiotic conditions are often expected to be less vulnerable to future increases in climatic variability. However, without defining the link between abiotic variability and the capacity for populations to resist and recover following disturbances (*i.e*., their resilience), we cannot predict the consequences of ongoing community reassembly. Evaluating the association between measures of abiotic variability and the resilience attributes of 2,242 animal, plant, and algae populations, we discredit the assumption that contemporary exposure to more frequent environmental shifts confers a greater ability to cope with future global change. Instead, the resilience attributes of natural populations appear to have been moulded over longer-term evolutionary timeframes and are thus not a response to more recent experiences.

## Introduction

Ongoing global shifts in the timing and magnitude of environmental variation warrant an understanding for the processes underlying the capacity for populations to resist and recover from disturbances (1, 2) (*i.e*., their resilience [3]). However, half a century since Holling first defined resilience in ecological systems (3), we still do not know whether and how past environmental regimes shape the resilience of extant species (4). Resolving this knowledge gap is pivotal for identifying those species most vulnerable to future increases in environmental stochasticity (5, 6), and thus for designing effective ecosystem management strategies (7).

While the resilience of ecological systems has attracted much attention for decades (8, 9), approaches to evaluate the resilience of natural populations and communities often overlook its short-term nature (10, 11) (but see (12)). Classic life-history theory posits that organisms operate under a strong trade-off coordinating their investments across different vital rates (13, 14). Indeed, considerable attention has been directed at using the vital rates of survival, progression (*e.g*., growth, development), retrogression (*e.g*., shrinkage [15], rejuvenation [16]), and reproduction to describe how energetic trade-offs shape population performance when exposed to disturbances (17–21) (*i.e*., external (a)biotic forces that can modify the composition of a system). Yet, attention has largely focused on the use of stochastic derivatives of long-term (*i.e*., asymptotic) population characteristics (such as stochastic population growth rate, *λ_s_*) as measures of population performance in the face of recurrent disturbances (22). However, disturbances can and oftentimes displace populations into a transient (*i.e*., short-term) phase, where their dynamics can vary considerably from their asymptotic trajectories (23) (Fig. 1A). Long-term measures of population performance are, thus, unlikely to reveal how environmental stochasticity influences the resilience of populations (10). Instead, transient population metrics, describing how the dynamics of populations can change following disturbance relative to their long-term characteristics (24, 25), can offer greater insight.

**Figure 1.**
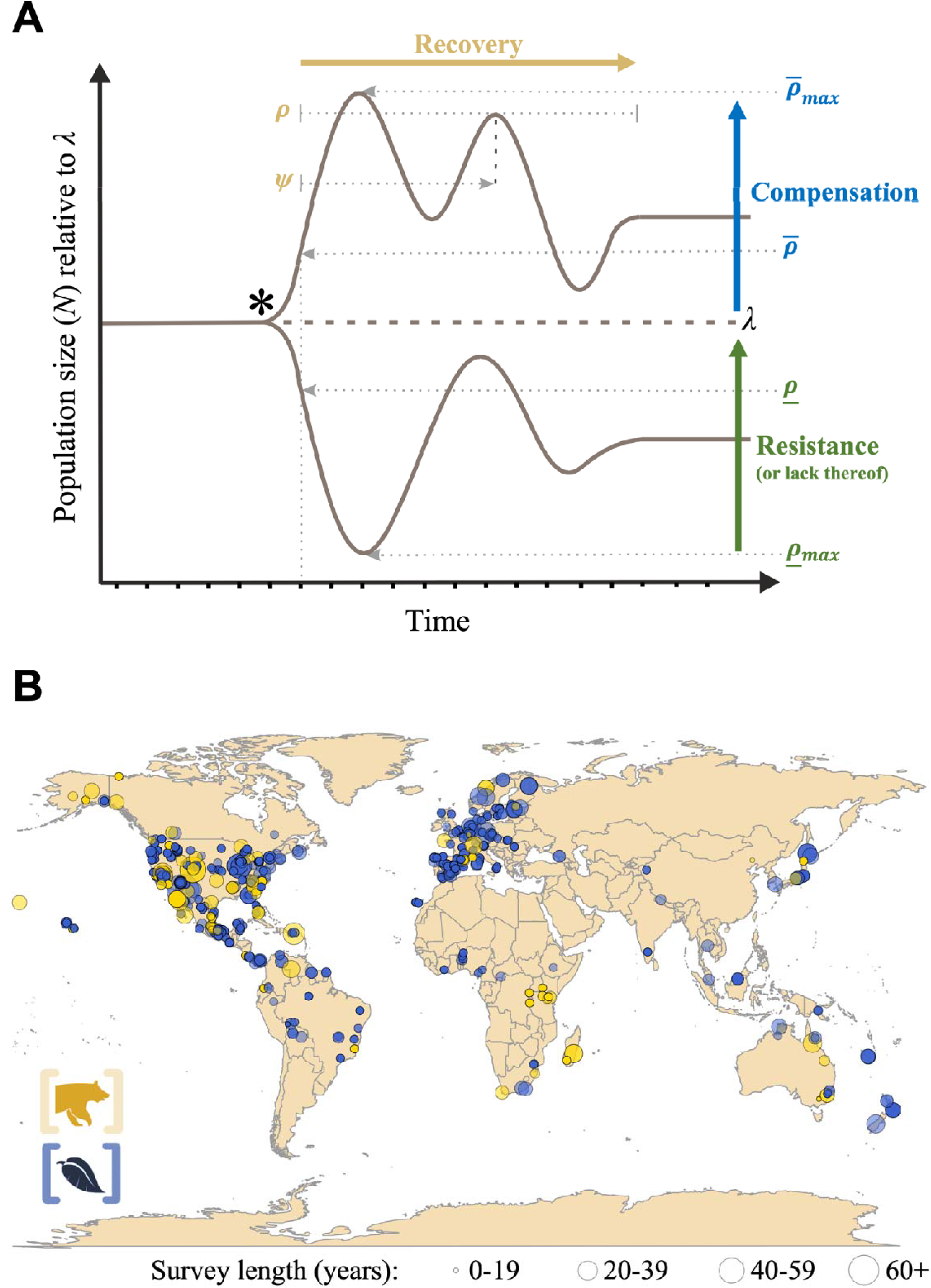
Demographic resistance, compensation, and time of recovery across 2,242 natural populations show how population resilience corresponds with exposure to environmental stochasticity. (**A**) Within stochastic environments, the dynamics of populations can vary considerably from long-term (*i.e*., asymptotic) expectations. Under stable stationary conditions, populations display asymptotic growth trajectories whereby the size of a population changes at a constant rate (*λ*). However, following a disturbance (*), populations can enter a transient state during which their growth rate can differ significantly from asymptotic expectations. The duration and form of this transient phase depends, in addition to key moments of disturbances (*e.g*., magnitude, frequency, *etc*.), on a population’s resilience attributes of resistance, compensation, and time to recovery. Here, *resistance* is the ability for a population to avoid a decline in size following a disturbance, whereas *compensation* is the extent to which a population may increase in size following a disturbance. Meanwhile, time of recovery (henceforth *recovery*) is the duration needed for a population to converge back to its stationary equilibrium following a disturbance. Transient demographic theory offers the opportunity to quantify the short-term dynamics of natural populations, thus unlocking the potential for macroecological studies exploring patterns and plausible mechanisms of demographic resilience. Transient increases in population size (*N*) can be evaluated using metrics of population reactivity (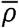; increase in *N* within a one-time step following a disturbance) and maximal amplification (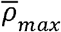; maximum increase in *N* during the transient period). Equally, the magnitude of transient declines in population size can be assessed using the metrics of first-step attenuation (*<u>ρ</u>*; decrease in *N* within a one-time step following a disturbance) and maximal attenuation (*<u>ρ</u>_max_*; maximum decrease in *N* during transient period). Finally, the damping ratio (*ρ*; rate of convergence back to stationary stability), and period of oscillation (*ψ*; time between corresponding phases of the largest oscillatory cycle in *N*) of populations offer insights into their capacity to go back to a stationary stable equilibrium. (**B**) We quantified the aforementioned transient metrics of demographic resilience for 2,242 populations from 369 species for whom Matrix Population Models (MPMs) are available from the global COMPADRE^25^ (Blue; plants & algae) and COMADRE^24^ databases (Orange; animals). Point size illustrates the relative length of time over which each population was surveyed to parameterise the MPMs deposited within COMPADRE and COMADRE.

Natural populations exposed to frequent disturbances are expected to experience a selection for traits, and vital rates, that enhance their resilience (26, 27). Within more stochastic environments populations endure more frequent (a)biotic fluctuations, each with the capacity to impose shifts in their structural composition. Following these disturbances, characteristics that inhibit transient declines in population growth, or those promoting short-term increases are likely more beneficial than those maximising long-term performance (12, 28). Logically, therefore, increased stochasticity should select for increased *resistance (i.e*., a population’s ability to avoid a decline in size after disturbance) and/or *compensation* (the ability to increase in size after disturbance), and shorter *recovery time* (henceforth recovery, the time needed for converging back to a stationary equilibrium after disturbance) (29, 30) (Fig. 1A). Accordingly, evaluating variation in the transient characteristics of populations across a gradient in environmental stochasticity presents a unique opportunity for exploring how population resilience is shaped by stochastic environments (11, 29–31). Moreover, this approach represents a space-for-time substitute (32) for predicting the performance of populations under future scenarios of increased stochasticity and more frequent recurrent disturbances.

Here, we provide a global assessment of the environmental determinants of population resilience, examining how short-term environmental drivers mediate long-term evolutionary outputs. Specifically, we quantify the inherent capacity for resistance, compensation, and recovery of 2,242 natural populations across 61 animal, 305 plant, and 3 algae species from two global demographic databases (33, 34) (Fig. 1B & Supplementary Table S1). Building on the expectation that past disturbance regimes have shaped current resilience (27, 35), we anticipated that (H1a) exposure to higher frequency environmental stochasticity will select for faster recovery, whilst (H1b) exposure to broader spectra in environmental stochasticity selects for increased resistance and reduced compensation. With the vital rates of survival, progression, retrogression, and reproduction describing how trade-offs in individual-level fitness translates into population-level performance (36), we expected (H2) patterns in population resilience to correspond with the underlying vital rates of populations; with investments into individual survival associated with enhanced resistance and greater reproductive investments aligned with faster recovery.

## Results and Discussion

Sourcing climate records at the necessary temporal and spatial resolution for quantifying deep-time environmental legacies presents a considerable challenge (37, 38). Thus, a population’s resilience to future global change is more often inferred from its exposure to more contemporary conditions (4, 39). Here, however, using phylogenetically-corrected partial least squares regression (pPLS) and Pearson’s tests of correlation, we found that no measure of environmental stochasticity constitutes a strong predictor of variation in the resistance, compensation, and recovery attributes of populations (Fig. 2 & Table 1). This finding was insensitive to both phylogenetic imputation (40) (Fig. S3) and the length of time used to quantify recent-past exposure to environmental stochasticity (Fig. S6 & S7).

**Figure 2.**
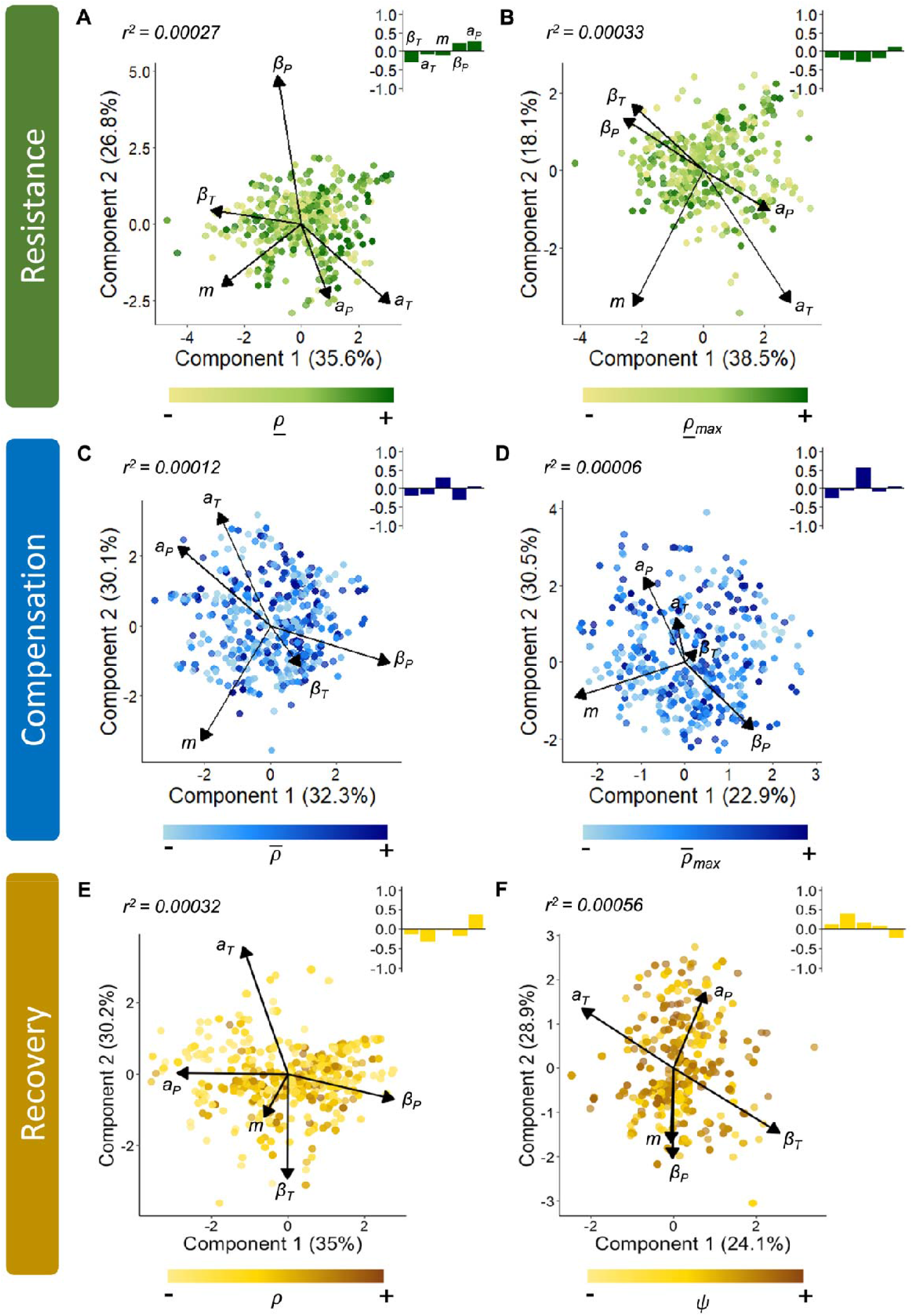
Variation across metrics of demographic resilience: resistance (green), compensation (blue), and recovery (orange) does not correspond with patterns in the exposure of populations to environmental stochasticity. Scores and loadings of a phylogenetically-weighted Partial Least Squares (pPLS) regression analysis exploring the correlation between patterns in the variation of the six transient metrics of (**A**) first-step attenuation (*<u>*ρ*</u>*), (**B**) maximal attenuation (*<u>ρ</u>_max_*), (**C**) reactivity 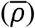, (**D**) maximal amplification 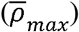, (**E**) damping ratio (*ρ*), and (**F**) period of oscillation (*ψ*), and the five metrics of environmental stochasticity: temperature frequency spectrum (*β_T_*), temperature autocorrelation (*a_T_*), thermal range/magnitude (*m*), precipitation frequency spectrum (*β_P_*), and precipitation autocorrelation (*a_P_*). The component scores along each axis display the percentage variance in the environmental stochasticity variables captured by each component, with the first two components alone explaining >50% of the variance across all models. The gradation in point colour then reflects patterns in the relative magnitude of each transient metric recorded from each population, with darker shades indicating higher estimates. Insert barplots are the standardised regression coefficients (*b*) highlighting the relative weighting of each abiotic variable in the overall capacity of each pPLS model to explain variation in a given transient metric (*r^2^*).

**Table 1.** The resilience attributes of resistance (green), compensation (blue), and recovery (orange) of natural populations do not correlate with their relative exposure to environmental stochasticity. Using phylogenetically-corrected Pearson’s tests of correlation, we explored the association between transient metrics of demographic resistance (first-timestep attenuation,_ & and maximal attenuation, __*max*_), compensation (reactivity,^−^ & maximal amplification, ^−^_*max*_), and recovery (damping ratio, *ρ* & period of oscillation, *ψ*; Fig. 1A) and five metrics of environmental stochasticity: temperature frequency spectrum (*β_T_*), temperature autocorrelation (*a_T_*), thermal range (*m*), precipitation frequency spectrum (*β_P_*), and precipitation autocorrelation (*a_P_*). Correlation displayed using Pearson’s correlation coefficient (*r*).

| Transient metric | Resilience attribute | $\beta_T$ | $a_T$ | $m$ | $\beta_P$ | $a_P$ |
| --- | --- | --- | --- | --- | --- | --- |
| $\underline{\rho}$ | Resistance | -0.0106 | 0.0060 | -0.0094 | -0.0024 | 0.0042 |
| $\underline{\rho}_{max}$ | | -0.0071 | 0.0031 | -0.0121 | -0.0093 | 0.0068 |
| $\bar{\rho}$ | Compensation | -0.0026 | 0.0007 | 0.0070 | -0.0080 | 0.0041 |
| $\bar{\rho}_{max}$ | | -0.0016 | <0.0001 | 0.0069 | -0.0026 | 0.0009 |
| $\rho$ | Recovery | 0.0012 | -0.0007 | 0.0026 | -0.0124 | 0.0147 |
| $\psi$ | | -0.0071 | 0.0133 | 0.0052 | 0.0037 | -0.0070 |

Using a novel framework, based on punctual disturbances altering population structure, to quantify demographic resilience (30), we estimated the six transient metrics comprising resistance (first-step attenuation, *<u>ρ</u>* & maximal attenuation, *<u>ρ</u>_max_*), compensation (reactivity, 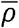 & maximal amplification, 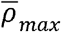), and recovery (damping ratio, *ρ* & period of oscillation, *ψ*) in each population following disturbance (25, 30, 41) (Fig. 1A). Next, we quantified the exposure of these populations to environmental stochasticity using measures of mean thermal range (*m*), as well as the spectral frequency and autocorrelation of temperature (*β_T_* & *a_T_*) and precipitation regimes (*β_P_* & *a_P_*) they experienced during the 50-years preceding the beginning of each study (Supplementary S4). Over any given timeframe, shorter-lived populations (mean life expectancy ≤ 10 years) are likely to have had a larger number of generations exposed to local stochasticity, thereby offering greater opportunity for adaptive change. Yet, our finding persists irrespective of mean life expectancy, with recent-past exposure to environmental stochasticity having an equally negligible influence on the resilience attributes of populations of both long- and short-lived species (Fig. S5).

Resistance and compensation, but not recovery, are determined by energetic investments across somatic maintenance/development and reproduction. Using pPLS across our 2,242 populations, we evaluated the relationship between our six measures of demographic resilience (Fig. 1A) and their sensitivities to each of the vital rates of survival (*σ*), progression (*γ*), retrogression (*τ*), and fecundity (*φ*; Fig. 3). These vital rate sensitivities reflect how much each transient metric would change following perturbations in each vital rate (41). Thus, these sensitivities highlight how investments into any one vital rate influence a populations’ capacity to resist, compensate, and recover following a disturbance, and so provide a measure of the absolute importance of each vital rate in shaping demographic resilience. We focused on sensitivities here, rather than elasticities (proportional sensitivities [42]), as the former provide a closer representation of selection gradients (43). Patterns across the vital rate sensitivities of each transient characteristic corroborate recent evidence that the resistance and compensation attributes of populations are constrained by individual-level investments across survival, progression, retrogression, and reproduction (44). Meanwhile, we found no evidence of demographic selection pressures on the attribute of recovery (Fig. 3 & Table 2). Again, these findings are insensitive to phylogenetic imputations (Fig. S4).

**Figure 3.**
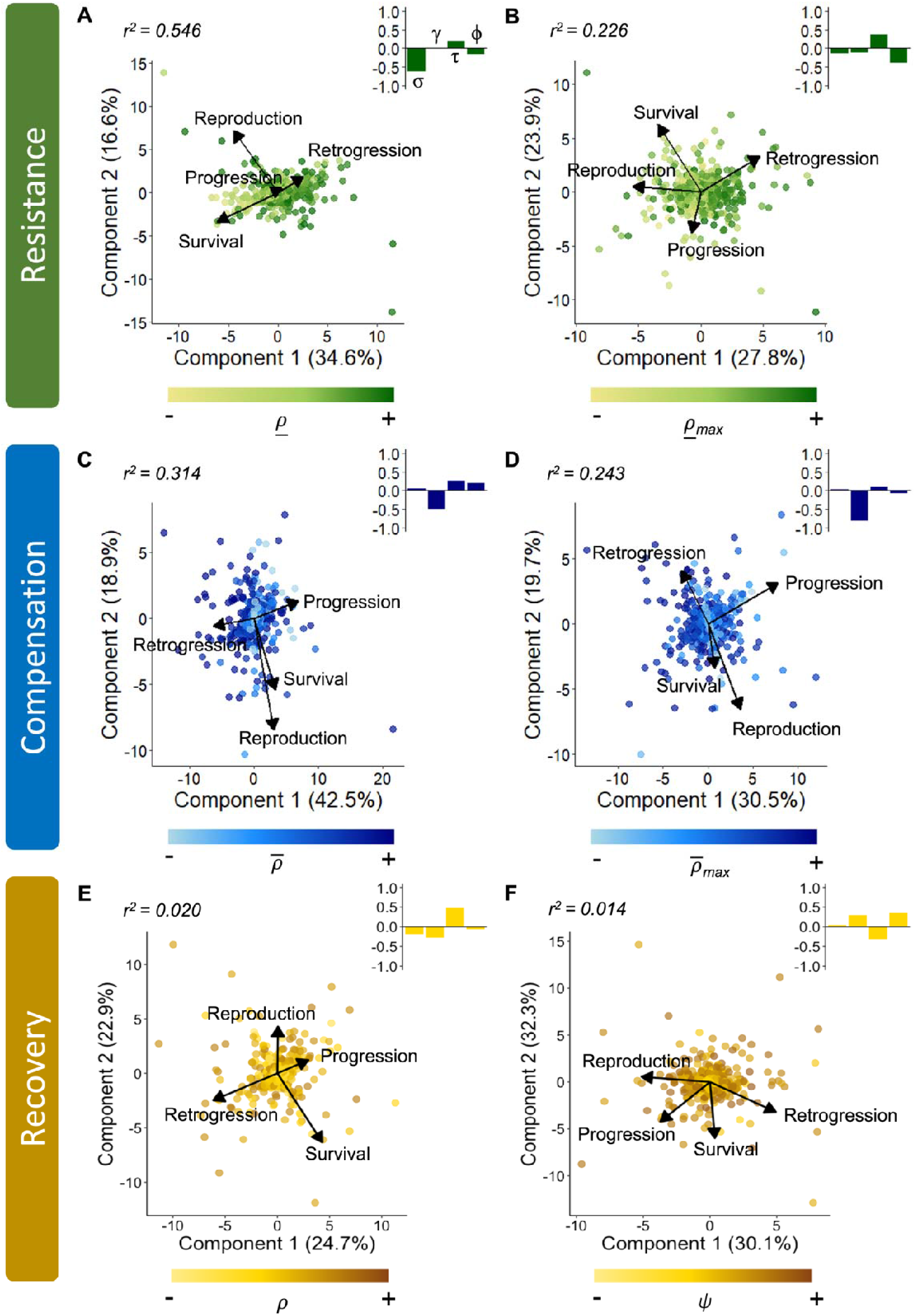
The resilience attributes of resistance (green), compensation (blue), and recovery (orange), in natural populations are determined by the relative energetic investments of their individuals. Scores and loadings of a phylogenetically weighted Partial Least Squares regression analysis exploring the sensitivity patterns of the six transient metrics of (**A**) first-step attenuation (<u>*ρ*</u>), (**B**) maximal attenuation (*<u>ρ</u>_max_*), (**C**) reactivity 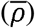, (**D**) maximal amplification 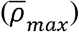, (**E**) damping ratio (*ρ*), and (**F**) period of oscillation (*ψ*), towards the vital rates of survival (*σ*), progression (*σ*), retrogression (*τ*), and reproduction (□). The component scores along each axis display the percentage variance in the environmental stochasticity variables captured by each component, with the first two components alone explaining >47% of the variance across all models. The gradation in point colour then reflects patterns in the relative magnitude of each transient metric recorded from each population, with darker shades indicating higher estimates. Insert barplots are the standardised regression coefficients (*b*) highlighting the relative weighting of each vital rate in the overall capacity of each pPLS model to explain variation in a given transient metric (*r^2^*).

**Table 2.** Variation across resistance (green) and compensation (blue) of natural populations corresponds with the energetic investments of their individuals, but characteristics of recovery (orange) do not. Using phylogenetically-corrected Pearson’s tests of correlation, we explored the association between the transient metrics of resistance (first-timestep attenuation,_ & and maximal attenuation, __*max*_), compensation (reactivity,^−^ & maximal amplification, ^−^_*max*_), and recovery (damping ratio, *ρ* & period of oscillation, *ψ*; Fig. 1A) and their sensitivities to the vital rates of survival (*σ*), progression (*σ*), retrogression (*τ*), and reproduction (□). Correlation displayed using Pearson’s correlation coefficient (*r*), with the colour gradation highlighting the strength of the absolute correlation between each transient metric and its vital rate sensitivities, with darker shades representing stronger associations.

| **Transient metric** | **Resilience attribute** | *Survival*<br>$\sigma$ | *Progression*<br>$\gamma$ | *Retrogression*<br>$\tau$ | *Reproduction*<br>$\phi$ |
| --- | --- | --- | --- | --- | --- |
| $\rho_{-}$ | Resistance | -0.70 | -0.07 | 0.28 | -0.39 |
| $\rho_{-max}$ | | -0.17 | 0.07 | 0.31 | -0.34 |
| $\bar{\rho}$ | Compensation | -0.11 | -0.49 | 0.39 | -0.04 |
| $\bar{\rho}_{max}$ | | -0.02 | -0.49 | -0.13 | -0.14 |
| $\rho$ | Recovery | -0.06 | -0.06 | 0.11 | -0.01 |
| $\psi$ | | <0.01 | 0.06 | -0.08 | 0.08 |

A strong phylogenetic signal across energetic investments into resistance, compensation, and recovery suggests that species demographic resilience is not shaped by recent environmental patterns, but by phylogenetic ancestry. Using estimates of phylogenetic signal (Pagel’s *λ* [45]; not to be confused with long-term population growth rate, *λ*) we demonstrate a strong phylogenetic signal (Pagel’s *λ* > 0.940, p <0.001) across the transient metrics of resistance (first-timestep attenuation & maximal attenuation) and compensation (reactivity & maximal amplification), and their corresponding vital rate sensitivities (Table 3). Phylogenetic signal reflects the proportion of variation in a trait across different species that can be explained by their shared evolutionary history (45, 46). As such, this analysis quantifies the extent to which evolution may have shaped the observed resilience attributes of natural populations. We observed little association between the phylogenetic patterns of resilience and the environmental stochasticity regimes to which our examined natural populations were exposed. Selection pressures constrain how individuals allocate finite resources across survival, somatic development, and reproduction, thus mediating the capacity for populations to exploit and prevail within their local environments (14, 26). Over time, these selective forces have moulded the resilience attributes of populations, but not in response to the stochasticity regimes they currently endure (47, 48). It is necessary, therefore, to highlight the limitations associated with using the recent exposure of populations to environmental stochasticity as a predictor of their continued resilience to future disturbances.

**Table 3.**
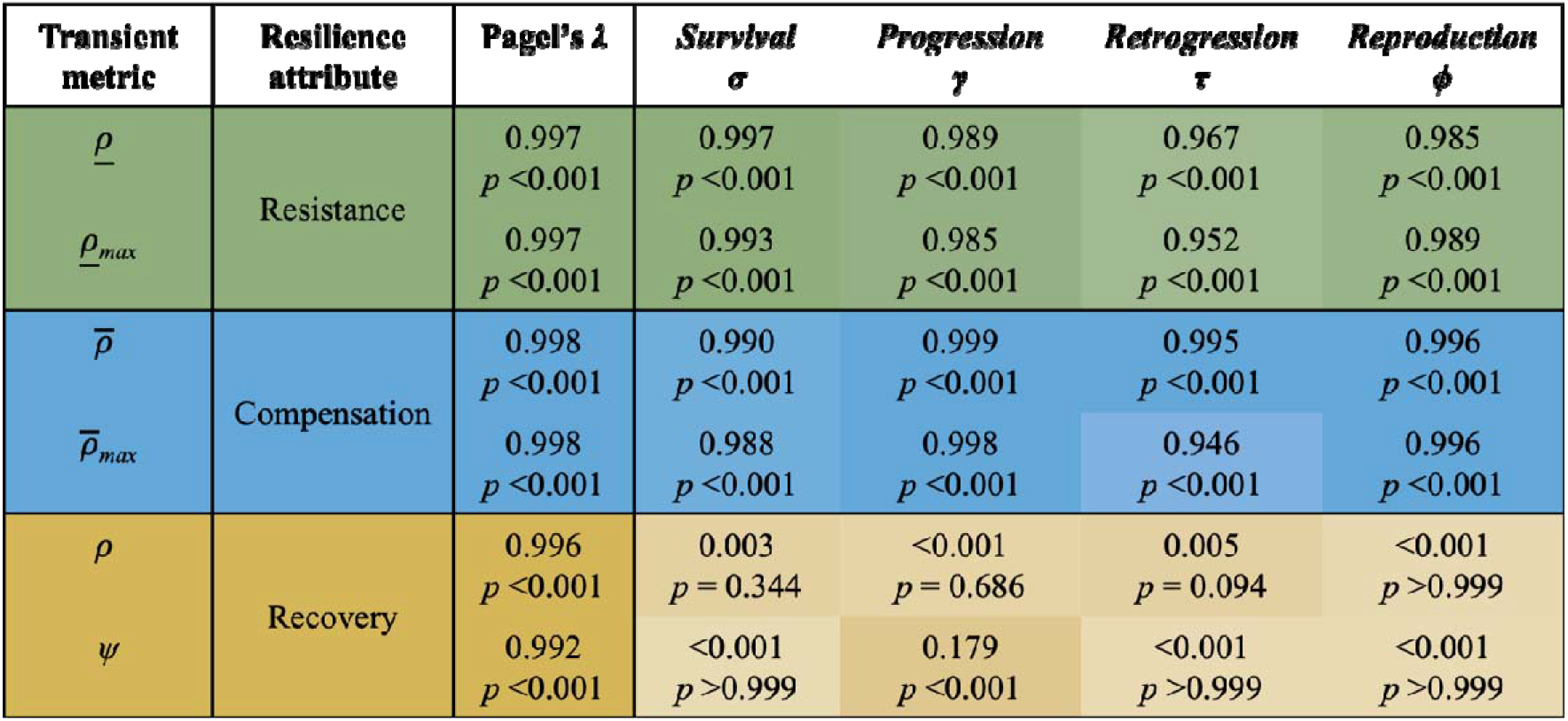
Variation in resistance (green), compensation (blue), and recovery (orange) is strongly predicted by phylogenetic association. However, whilst a strong phylogenetic signal is also evident across the vital rate sensitivities of measures of resistance and compensation, there is a negligible phylogenetic signal across the vital rate sensitivities of measures of recovery. To quantify the strength of statistical non-independence in the resilience attributes of natural populations due to species’ common ancestry, we estimated the phylogenetic signal (Pagel’s λ(45)) across our transient metrics of demographic resistance (first-timestep attenuation, _ & and maximal attenuation, __*max*_), compensation (reactivity,^−^ & maximal amplification, ^−^_*max*_), and recovery (damping ratio, *ρ* & period of oscillation, *ψ*; Fig. 1A), as well as their sensitivities to the vital rates of survival (*σ*), progression (*σ*), retrogression (*τ*), and reproduction (*□*). Pagel’s *λ*(45) ranges between 0, indicating that traits have evolved independently of phylogeny, and 1, representing a high phylogenetic signal. Colour gradation highlights the likelihood that the phylogenetic signal observed across each transient metric and its vital rate sensitivities differs significantly from 0 (*p* <0.05), with darker shades representing stronger signals.

Caution is necessary when interpreting our results regarding the selection pressures maintained by environmental stochasticity. Our exploration into the environmental drivers of demographic resilience focuses only on terrestrial populations. Compared with marine systems, terrestrial environments afford organisms with more opportunities for seeking out tolerable microclimates (49), thereby reducing the susceptibility of terrestrial taxa to environmental shifts. By contrast, marine species inhabit conditions closer to their physiological limits (50), rendering them more sensitive to abiotic shifts. More importantly, though, here we quantify past exposure to environmental stochasticity using maximum exposure legacies of 100-years. It is plausible, however, that deep-time environmental regimes would offer greater predictive potential. Indeed, deep-time studies have demonstrated how the latitudinal diversity gradient, a prominent pattern underlying contemporary ecological understanding, has only persisted during the past 30 million years (51). Whilst environmental stochasticity is known to influence population dynamics (52), its observable effects on population characteristics can remain negligible until compounded by external factors such as changing habitat configurations (53). Any direct impacts of stochasticity on the resilience of natural populations may, therefore, become more detectable with time.

Despite the potential influence of historical climate legacies on the resilience of natural populations, our results show a robust lack of influence of recent-past climate on population resilience, even after explicitly accounting for lifespan and legacy time. Adaptation is the accumulation of beneficial, heritable characteristics within a population over multiple generations. Therefore, populations exposed to specific conditions over extended timeframes, or those that turn over multiple generations within a short period of time, are more likely to display trait characteristics adapted to those conditions (47, 48, 54). To evaluate the implications of differing exposure periods on our observations, we first repeated our pPLS analyses exploring the relationship between demographic resilience and environmental stochasticity focusing on only populations of short-lived species (Supplementary S6). We determined population longevity using estimates of mean life expectancy (*η_e_*), with populations of *η_e_* ≤10-years defined as short-lived (n = 1,606 populations). Next, we repeated our analyses using only populations for which 100-year abiotic legacies could be sourced (Supplementary S7). Neither approach improved our ability to predict the demographic resilience attributes of natural populations using measures of environmental stochasticity (Figs. S5, S6 & S7).

### Conclusions

Global change is changing the periodicity of phenological drivers (55, 56), reducing return times between severe disturbance events (57, 58). Natural populations worldwide are thus being exposed to increasingly stochastic environments, with many facing imminent collapse(59). Here, we illustrate, however, that the recent-past exposure (last 50-100 years) of natural populations to environmental stochasticity does not constrain, nor guarantee, their resilience towards future climatic shifts. This realisation that population resilience appears constrained by the adaptation of species over longer-term evolutionary timeframes, not the conditions experienced by populations following their dispersal, is not trivial. Rather, this key insight can help to focus current debates on the concept of resilience (9, 29, 60), enabling our attention to be directed towards identifying populations with high adaptive potential, and maintaining the high genetic diversity necessary to enhance the resilience of natural populations. Reliant on preserving and enhancing the earth’s natural capital, initiatives such as the United Nations Sustainable Development Agenda are interwoven with the conservation-orientated goals of our responsibility for environmental stewardship (61, 62). Accordingly, a detailed understanding of the mechanisms that do and do not promote the resilience of natural populations is imperative for not only achieving global conservation success, but also for securing our socioeconomic future.

## Methods

### Demographic data extraction & transient indices

To evaluate the selection pressures on the resilience attributes of natural populations, we extracted Matrix Population Models (MPMs) from the open-source COMPADRE Plant Matrix Database (34) (v. 5.0.1) and COMADRE Animal Matrix Database (33) (v. 3.0.1). MPMs are discrete time, structured population models where the lifecycle is categorised into discrete state classes (*i.e*., age, size, and/or developmental stages) (41). Combined, COMPADRE and COMADRE contain over 12,000 MPMs from more than 1,100 animal and plant species. However, here we only retained MPMs satisfying the following six criteria to test our hypotheses: 1) MPMs reflecting the demographic characteristics of individual populations recorded across a single time period (*Individual* MPMs). We did, however, select MPMs consisting of demographic information averaged across multiple populations and/or time periods (*mean* MPMs), for populations for which no individual matrices were available (330 populations after applying the additional criteria below); 2) MPMs based on annual surveys to ensure all subsequent metrics obtained reflected identical units of time, thus allowing for their comparability; 3) MPMs representing wild, un-manipulated populations, to guarantee investigating the selection pressures underpinning the resilience of natural populations and the possibility to link their dynamics to their local environmental regimes; 4) MPMs comprised of three or more life stages, as lower dimension MPMs typically lack the necessary resolution for estimating vital rates (63) and transient dynamics (64); 5) MPMs from populations with known latitude and longitude information to allow us to link their demographic properties to local environmental regimes; and finally 6) MPMs describing full life cycles (e.g., no missing data on survival, progression, retrogression, and reproduction) to ensure the correct calculation of vital rates and transient metrics. Following these criteria, we retained 3,890 MPMs corresponding with 3,204 populations across 438 plant species, 665 populations across 112 animal species, and 21 populations across six algae species (Supplementary Table S1).

We further refined our list of MPMs according to their transient, asymptotic, and species-specific properties. All MPMs were tested for irreducibility (*i.e*., all life cycle stages are either directly, or indirectly connected to one another), ergodicity (*i.e*., asymptotic dynamics are independent of the initial population structure), and primitivity (*i.e*., MPMs consist of non-negative elements) (41). We carried out these tests using the corresponding *isErgodic, isIrreducible*, and *isPrimitive*, diagnostic functions from the R package *‘popdemo’* (65). A total of 1,203 reducible, imprimitive, and/or non-ergodic MPMs were excluded from further analyses on the basis that they represent untenable life cycles that defy logical biological processes (66). MPMs with population growth rates *λ*>2, indicating that the population is projected to increase two-fold or more every year, were also rejected as they represent unlikely realisations of population performance in our experience. Equally, MPMs from highly migratory (e.g., > 1,000 km) species were discarded, since their vital rate schedules are unlikely to be mostly shaped by the environment in which they were measured. We also note here that, across our initial population sample, the vital rate of clonality (*κ*) was rare, with only 140 populations across 37 plant species, and two populations from one animal species (*Amphimedon compressa* [67]) explicitly exhibiting this demographic process. Thus, to focus our analyses on common demographic currencies, we excluded all populations exhibiting clonality. Overall, our strict selection criteria resulted in a final sample of 2,242 MPMs, corresponding with 369 species: 402 populations from 61 animal species, 1,830 populations from 305 plant species, and 10 populations from three species of algae (Supplementary Table S1).

For each retained MPM, we calculated six transient metrics quantifying each population’s potential for demographic resistance (first-timestep attenuation, *<u>ρ</u>* & and maximal attenuation, *ρ_max_*), compensation (reactivity, 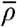 & maximal amplification, 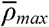), and recovery (damping ratio, *ρ* & period of oscillation, *ψ*), following a disturbance (30) (Fig. 1). Firstly, with estimates of transient dynamics known to be contingent on the reproductive strategies of populations, it was necessary to convert all post-reproductive matrices into a pre-reproductive format by adjusting patterns of reproduction to include a measure of adult survival (68). All MPMs were then standardised to separate their transient and asymptotic properties by dividing each matrix element by the MPM’s dominant eigenvalue, *λ* (24, 41). Following standardisation, from each MPM we obtained estimates of reactivity and first-timestep attenuation, as the upper and lower bounds returned by the function *reac*, and calculated maximal amplification and maximal attenuation indices using their respective functions *maxamp* and *maxatt*, all within the R package ‘*popdemo*’ (65). Meanwhile, we calculated the damping ratio of each MPM using *eigen.analysis* from the ‘*popbio*’ package (69). Finally, we calculated the period of oscillation (*ψ*) for each MPM using its subdominant eigenvalue (*λ*_2_) (equation 1) (41).

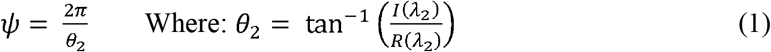

To explore how the fitness components of individuals mediate the selection gradients placed on demographic resilience by environmental variability, we calculated the sensitivity of each transient metric towards the vital rates of survival (*σ*), progression (*γ*), retrogression (*τ*), and fecundity (*φ*). For each MPM, we first estimated all vital rate sensitivities from their element-level constituents. Individual elements *a_ij_* within the MPM ***A*** typically describe combinations of multiple vital rates (70). Subsequently, calculating the sensitivity of each transient metric (*s_x_*) with respect to underlying vital rates requires the decomposition of element-level sensitivities into their vital rate components (70). Briefly, this decomposition requires the estimation of stage-specific survival probabilities (*σ_j_*) for each MPM. These estimates of *σ_j_*, are then used to determine the proportion of each matrix element *a_ij_* corresponding with survival (*σ*), progression (*γ*), retrogression (*τ*), and fecundity (*φ*) (70). We initially calculated the sensitivity of each transient metric at the matrix element-level (*s_ij_*). The sensitivities of the damping ratio (*ρ*) and period of oscillation (*ψ*) were determined using equations 2 & 3, with 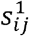 and 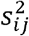 as the matrix element-level sensitivity matrices of the dominant and subdominant eigenvalues, respectively (41).

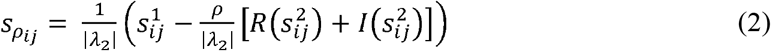

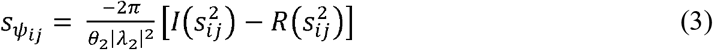

The sensitivities of reactivity, amplification, and attenuation (*s_ij_*) with respect to element *a_ij_* were then estimated as the magnitude of change (*δ*) in each transient metric (*x*) following a small change (here 0.01) in *a_ij_* (Equation 4) (71).

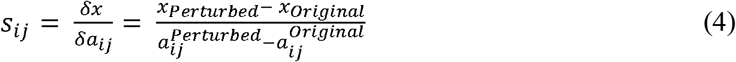

The distributions of each transient metric and its corresponding vital rate sensitivities were checked prior to subsequent regression analyses. To ensure normality across each distribution, outliers, defined as values outside the 95% confidence intervals of the distribution, were omitted, with the remaining estimates then transformed if necessary. For each transient metric, power transformations (*y^x^*) were used to achieve approximate normality using the Box-Cox transformation functions of the R package ‘*caret*’ (72) to estimate *x*. The distributions of damping ratio, period of oscillation, reactivity, and maximal amplification raised negative *x* values, and so their transformations took the form *1/y^|x|^*. Inverse and log transformations were also necessary for several of the vital rate sensitivity variables (See Table S2 for further details).

### Phylogenetic correction

Evaluating the selection pressures exerted on attributes of demographic resilience across multiple species requires an explicit consideration for how traits are expected to covary due to phylogenetic relationships (46, 73, 74). To account for such relationships in our analyses, we constructed a population-within-species level phylogenetic tree using phylogenetic data extracted from the Open tree of Life for the animal, plant, and algae species across our population sample (OTL [75]; Supplementary S3). Our approach here also allowed us to accommodate studies that included multiple, separate populations for the same species (*see below*). Firstly, the scientific names of each species associated with our extracted MPMs were checked against current taxonomy records using the R package *‘taxize’* (76). Next, we extracted information regarding the taxonomic classification and phylogeny of each species from the OTL database with the R package ‘*rotl*’ (77). Subsequently, using the ‘*ape*’ (78) and ‘*phytools*’ packages (79), this phylogenetic information was used to construct a species-level phylogenetic tree corresponding with the 369 unique species within our MPM list.

Beyond accounting for phylogenetic signals in trait variance-covariance across our population sample, it was necessary to ensure that our phylogenetic tree reflected the influence of spatial signals in the development of traits within species. Thus, we expanded our phylogenetic tree by adding branch tips to incorporate multiple population entries per species (*sensu* [80]), generating a population-level tree comprising our full sample of 2,242 populations (Supplementary S3). Finally, we computed the branch lengths for our phylogenetic tree using the function *compute.brlen* in the R package ‘*ape*’ (78). These branch lengths were estimated using Grafen’s arbitrary branch lengths (81), assuming a Brownian motion model with the variance between species directly proportional to time since divergence (82). Importantly, we constrained branch lengths between populations of the same species to approximately zero (0.0000001) under the assumption of negligible phylogenetic distance between species replicate populations.

### Quantifying environmental variability

To investigate the role of environmental selection pressures on the resistance, compensation, and recovery components of resilience in natural populations, we used a pPLS regression exploring the association between transient characteristics and metrics of environmental variability. We quantified the magnitude and frequency of environmental variation to which each population was exposed using the geographical location information extracted with each MPM from COMPADRE & COMADRE (33, 34) (Supplementary S4). Since temperature and precipitation rates are universal drivers of biological community assembly across terrestrial environments (83), we selected data describing temporal trends in thermal and precipitation regimes as a measure of the environmental variability experienced by each population. Crucially, however, with precipitation not directly influencing marine environments (although see [84]), we excluded marine populations (29 populations from six animal species, and 10 populations from three algal species) from this portion of our analyses.

We quantified environmental variance through the metrics of autocorrelation, abiotic range, and frequency spectrum using long-term temperature and precipitation records sourced from the CHELSA climate database (85). For each population, we extracted monthly records of maximum and minimum temperatures (°C) and mean precipitation rates (kg m^−2^) corresponding with the specific period during which the population was surveyed as detailed in COMPADRE and COMADRE, plus an additional 50-years prior to the onset of censusing to account for environmental legacy effects (86). We condensed maximum and minimum temperature readings into monthly estimates of mean temperatures and thermal range. Next, as a gauge of disturbance magnitude (*m*), we estimated the mean thermal range experienced by each population across their associated temporal records. We then arranged our monthly mean temperature and precipitation estimates into time series depicting the temporal environmental regimes to which each population was exposed. Using the ‘*colorednoise*’ package (87), we calculated the autocorrelation of each temperature (*a_T_*) and precipitation (*a_P_*) time series as a measure of environmental predictability (88).

The colour of environmental variation is depicted on a red to blue colour scale, from lower to higher frequencies, respectively (88). The frequency spectrum of a time series is expressed by its spectral exponent (*β*), which is calculated as the negative slope of the linear regression between the log spectral density and log frequency of the time series (89). We calculated the frequency spectrum of each precipitation (*β_P_*) and temperature (*β_T_*) time series as an indicator of the colour of environmental variation experience by each population (89). The spectral exponent for each time series was estimated using the spectrum function from the R package ‘*stats*’ (90).

### Partial Least Squares Regression

We utilised a phylogenetically corrected Partial Least Squares regression (pPLS) framework to test our hypothesis that the resistance, compensatory, and recovery of natural populations correspond with gradients in environmental variability and evaluate how this is mediated by fitness investments. Using a pPLS, we evaluated the relationship between our estimates of population resilience, and both associated environmental variability regimes and their vital rate sensitivities. The pPLS technique is considered a more powerful comparative tool than other available multivariate regression methods (91), as it simultaneously condenses the variation among numerous predictors whilst maximising the variance explained among response variables. Subsequently, we investigated the selection pressures on the resistance, compensation, and recovery attributes of natural populations, and therefore the capacity for environmental legacies, and vital rate characteristics, to serve as predictors of resilience attributes.

We first applied a phylogenetically-corrected Pearson’s test of correlation and pPLS to analyse the correlation between environmental variability and transient demographic characteristics. This approach enabled us to test for covariation between the transient characteristics of populations and gradients in their exposure to environmental variability. pPLS tests were carried out for each transient measure with the predictor variable set comprised of our five metrics of environmental variability. From each test, we then extracted component scores and loadings, which describe the arrangement of the environmental predictor variables within a multivariate space. We also obtained the percentage variance (%*var*) among the predictors explained by each regression component and the proportion of variance in the transient response variable explained by each component (*r^2^*) to estimate the strength of any association between environmental variability and the transient dynamics of our population sample.

Phylogenetically corrected correlation tests and pPLS analyses were used to examine for patterns between each transient characteristics and its associated vital rate sensitivities. Again, test coefficients (*r*), component scores, loadings, %*var*, and *r^2^* values were calculated to quantify the influence of the fitness components of survival, progression, retrogression, and reproduction towards the transient characteristics of natural populations. All pPLS analyses were conducted using the ‘*pls*’ R package (92), with modifications included to ensure our analyses accounted for any evolutionary covariance between the sensitivity patterns and transient characteristics of different populations (79, 93, 94).

We carried out all pPLS analyses using only complete entries, omitting populations missing estimates for any one variable. To provide further clarity regarding any patterns we observed between our measures of population resilience, their environmental legacies, and their vital-rate sensitivities, we repeated each analysis twice. During these repeated tests, we first evaluated whether considering the life expectancies of populations influenced any observed patterns (Supplementary S6). Within a given time period, long-lived species will likely experience fewer generations than shorter-lived species diminishing the relative impact of existing selection pressures on their trait characteristics (95). Thus, it was necessary to ensure that the inclusion of long-lived species within our population sample did not limit our capacity for exploring environmental selection pressures. We categorised each population within our sample as either long- or short-lived according to their associated mean life expectancy (η_e_), calculated from each extracted MPM using the R package ‘*IPMpack*’ (96). Next, we repeated our pPLS analyses using only ‘short-lived’ populations for which η_e_ ≤ 10 years (n = 1606 populations). This threshold was selected as a balance between maximising the number of generations experienced by populations during the multi-year time series used in calculating environmental legacies and maximising our sample size.

Understanding patterns in the responses of natural populations to environmental change is often impeded by gaps in available data linking population characteristics to abiotic regimes (97). To ensure that missing data was not obscuring any correlation between our measures of population resilience and environmental stochasticity we repeated our analyses using phylogenetic imputation to estimate any demographic measures of resilience missing across our population sample (40) (Supplementary S5). We calculated the phylogenetic signal (Pagel’s *λ* [45]) of each transient and sensitivity variable using the *phylosig* function from the ‘*phytools*’ package (79). Pagel’s *λ* exists on the scale 0 < *λ* > 1, with 0 indicating traits have evolved independently of phylogeny, and 1 representing a high phylogenetic signal (45). For any variable exhibiting a strong phylogenetic signal (*i.e*., Pagel’s *λ* ≥ 0.65) that differs significantly from zero (*p* < 0.05), we imputed all missing values assuming a Brownian motion evolutionary model, before repeating our pPLS analyses.

## Acknowledgments

Funding for this research was provided by a DTP Natural Environment Research Council Scholarship to JC, a NERC IRF (NE/M018458/1) to RS-G, a Ramon Areces Foundation Postdoctoral Fellowship to PC, and Winifred Violet Scott Estate funding to MB. We acknowledge the University of Leeds High Performance Computing (ARC) team, who maintain the facilities used for the extraction of extensive climate records from CHELSA (https://chelsa-climate.org/).

## Author Contributions

JC, RS-G, MB, and PC conceived the ideas and methodology for the project. JC, PC, and RS-G carried out data extraction and analysis. JC led the writing of the manuscript, with close support from RS-G, and all authors contributing critically to the writing and giving final approval for publication.

## Data Accessibility

All Matrix Projection Models used within this study are available from the COMADRE and COMPADRE demographic databases (https://compadre-db.com/). Equally, all taxonomic information used in the construction of our phylogenetic trees can be sourced from the Open Tree of Life (https://tree.opentreeoflife.org/), and all temperature, and precipitation records used in this study can be downloaded from the CHELSA climate database (https://chelsa-climate.org/). Finally, the R code used for all the data extraction and analyses presented throughout this manuscript have been uploaded to GitHub and will be made publicly available following publication.

## Notes

### Competing Interest Statement

The authors have declared no competing interest.

### Summary of Updates

Shortened the manuscript to improve clarity

