## Supplementary material for "Recent exposure to environmental stochasticity does not determine the resilience of natural populations"

**S1. Species List**

**Table S1.** **A complete list of the 556 species for which data was extracted from the** **COMPADRE** (1) **& COMADRE** (2) **databases for this study. Entries are colour coded to distinguish each of the six algae (Grey), 112 animal (White), and 438 plant (Red) species extracted.** Each species is listed along with their corresponding realm classification (Terrestrial, Freshwater or Marine), the number of population replicates extracted initially, the number of population replicates retained in our final refined sample, and the original data source(s) (if provided). Population replicates dropped following initial extraction corresponded with either: (1) highly migratory species making it difficult to link demographic properties to the environment in which they were measured, or (2) Matrix Population Models (MPMs) exhibiting population growth rates >2, and/or (3) *reducible*, *non-ergodic* or *imprimitive* characteristics, and therefore representing untenable life cycles that defy logical biological processes.

|  | **Species** | **Realm** | **Extracted** | **Retained** | **Source** |
| --- | --- | --- | --- | --- | --- |
|  | *Ascophyllum nodosum* | Marine | 1 | 1 | (3, 4) |
|  | *Cystoseira zosteroides* | Marine | 4 |  | (5) |
|  | *Gracilaria gracilis* | Marine | 8 | 8 | (6) |
|  | *Laminaria digitata* | Marine | 1 | 1 | (7) |
|  | *Macrocystis pyrifera* | Marine | 1 |  | (8) |
|  | *Pterygophora californica* | Marine | 6 |  | (9) |
|  | *Alces alces* | Terrestrial | 15 | 8 | (10–12) |
|  | *Alouatta seniculus* | Terrestrial | 4 |  | (13) |
|  | *Amazona vittata* | Terrestrial | 1 | 1 | (14) |
|  | *Ambloplites rupestris* | Aquatic | 3 |  |  |
|  | *Ammocrypta pellucida* | Aquatic | 1 | 1 | (15) |
|  | *Amphimedon compressa* | Marine | 2 |  | (16) |
|  | *Anser anser* | Terrestrial | 1 |  | (17) |
|  | *Anthropoides paradiseus* | Terrestrial | 11 |  | (18) |
|  | *Astroblepus ubidiai* | Aquatic | 6 | 4 | (19) |
|  | *Bostrychia hagedash* | Terrestrial | 1 | 1 | (20) |
|  | *Brachyteles hypoxanthus* | Terrestrial | 25 | 25 | (21, 22) |
|  | *Buteo solitarius* | Terrestrial | 4 | 4 | (23) |
|  | *Calidris temminckii* | Terrestrial | 2 |  | (24) |
|  | *Callorhinus ursinus* | Marine | 1 |  | (25) |
|  | *Callospermophilus lateralis* | Terrestrial | 18 | 7 | (26) |
|  | *Calyptorhynchus lathami* | Terrestrial | 2 |  | (27) |
|  | *Canis lupus* | Terrestrial | 1 | 1 | (28) |
|  | *Capra ibex* | Terrestrial | 4 |  | (29) |
|  | *Cardisoma guanhumi* | Terrestrial | 3 | 3 |  |
|  | *Cebus capucinus* | Terrestrial | 22 | 22 | (21, 22) |
|  | *Cercopithecus mitis* | Terrestrial | 28 | 28 | (21) |
|  | *Certhia americana* | Terrestrial | 1 |  | (30) |
|  | *Cervus elaphus* | Terrestrial | 3 |  | (31–33) |
|  | *Chelodina expansa* | Aquatic | 2 | 2 | (34) |
|  | *Chelydra serpentina* | Aquatic | 1 | 1 | (35, 36) |
|  | *Chen caerulescens* | Terrestrial | 6 |  | (37) |
|  | *Chrysemys picta* | Aquatic | 3 | 1 | (38–40) |
|  | *Cicindela ohlone* | Terrestrial | 14 | 5 | (41) |
|  | *Colias alexandra* | Terrestrial | 5 |  | (42) |
|  | *Connochaetes taurinus* | Terrestrial | 2 |  | (43) |
|  | *Cottus bairdi* | Aquatic | 3 |  |  |
|  | *Crocodylus johnsoni* | Aquatic | 4 | 2 | (44) |
|  | *Crocodylus niloticus* | Aquatic | 2 |  | (45) |
|  | *Cryptophis nigrescens* | Terrestrial | 1 | 1 | (46) |
|  | *Dasypus novemcinctus* | Terrestrial | 3 | 3 | (47) |
|  | *Diadema antillarum* | Marine | 4 |  | (48) |
|  | *Dipodomys spectabilis* | terrestrial | 2 | 1 | (49) |
|  | *Eidolon helvum* | Terrestrial | 1 |  | (50) |
|  | *Elephas maximus* | Terrestrial | 1 |  | (51) |
|  | *Emydura macquarii* | Aquatic | 2 | 2 | (34) |
|  | *Epidalea calamita* | Aquatic | 1 |  | (52) |
|  | *Epinephelus morio* | Marine | 1 |  | (53) |
|  | *Etheostoma flabellare* | Aquatic | 3 |  |  |
|  | *Eumetopias jubatus* | Marine | 1 |  | (54) |
|  | *Falco peregrinus* | Terrestrial | 13 |  | (55) |
|  | *Fulmarus glacialis* | Terrestrial | 1 | 1 | (56) |
|  | *Giraffa camelopardalis* | Terrestrial | 4 | 4 | (57, 58) |
|  | *Gorilla beringei* | Terrestrial | 41 | 41 | (21, 22) |
|  | *Haliaeetus albicilla* | Terrestrial | 2 |  | (59) |
|  | *Homo sapiens* | Terrestrial | 26 |  |  |
|  | *Hoplocephalus bungaroides* | Terrestrial | 1 | 1 | (46) |
|  | *Hystrix refossa* | Terrestrial | 1 |  | (60) |
|  | *Kinosternon subrubrum* | Aquatic | 3 | 3 | (61) |
|  | *Kobus ellipsiprymnus defassa* | Terrestrial | 2 |  | (62) |
|  | *Lagopus muta* | Terrestrial | 1 | 1 | (63) |
|  | *Larus heermanni* | Terrestrial | 2 |  |  |
|  | *Leptonychotes weddellii* | Marine | 21 |  | (64) |
|  | *Macaca mulatta* | Terrestrial | 12 | 12 | (65–67) |
|  | *Malaclemys terrapin* | Aquatic | 1 | 1 | (68) |
|  | *Marmota flaviventris* | Terrestrial | 2 | 2 | (69) |
|  | *Montastraea annularis* | Marine | 13 |  | (70–72) |
|  | *Nipponia nippon* | Terrestrial | 1 |  | (73) |
|  | *Nocomis leptocephalus* | Aquatic | 3 |  |  |
|  | *Nuttallia obscurata* | Marine | 2 | 1 | (74) |
|  | *Odocoileus virginianus* | Terrestrial | 14 | 13 | (75) |
|  | *Oncorhynchus tshawytscha* | Marine | 5 |  | (76, 77) |
|  | *Onychogalea fraenata* | Terrestrial | 1 | 1 | (78) |
|  | *Ovis aries* | Terrestrial | 6 |  | (79) |
|  | *Ovis canadensis* | Terrestrial | 22 | 6 | (80–82) |
|  | *Pan troglodytes schweinfurthii* | Terrestrial | 45 | 45 | (21) |
|  | *Panthera pardus* | Terrestrial | 1 | 1 | (83) |
|  | *Papio cynocephalus* | Terrestrial | 37 | 37 | (21) |
|  | *Paramuricea clavata* | Marine | 11 | 10 | (84, 85) |
|  | *Phoca vitulina* | Marine | 1 |  | (86) |
|  | *Phoebastria immutabilis* | Terrestrial | 1 | 1 | (87) |
|  | *Phrynosoma cornutum* | Terrestrial | 2 | 2 | (88) |
|  | *Plexaura sp.* | Marine | 4 |  | (89) |
|  | *Pocillopora damicornis* | Marine | 1 |  | (90, 91) |
|  | *Podocnemis expansa* | Aquatic | 1 | 1 | (92) |
|  | *Presbytis thomasi* | Terrestrial | 1 |  | (93) |
|  | *Propithecus edwardsi* | Terrestrial | 2 | 2 | (94) |
|  | *Propithecus verreauxi* | Terrestrial | 24 | 24 | (21, 95) |
|  | *Rangifer tarandus platyrhynchus* | Terrestrial | 1 |  | (96) |
|  | *Rutilus rutilus* | Aquatic | 1 | 1 | (97) |
|  | *Saguinus fuscicollis* | Terrestrial | 4 |  | (98) |
|  | *Saguinus imperator* | Terrestrial | 3 |  | (98) |
|  | *Sceloporus grammicus* | Terrestrial | 8 | 8 | (99–101) |
|  | *Sceloporus mucronatus mucronatus* | Terrestrial | 1 | 1 | (102) |
|  | *Scolytus ventralis* | Terrestrial | 5 | 5 | (103) |
|  | *Spermophilus dauricus* | Terrestrial | 1 | 1 | (104) |
|  | *Sterna hirundo* | Terrestrial | 8 |  | (105) |
|  | *Sternotherus odoratus* | Aquatic | 2 |  | (40) |
|  | *Sternula antillarum browni* | Terrestrial | 1 |  |  |
|  | *Strix occidentalis occidentalis* | Terrestrial | 12 | 11 | (106) |
|  | *Sus scrofa scrofa* | Terrestrial | 1 |  | (107) |
|  | *Tamias striatus* | Terrestrial | 11 |  | (108) |
|  | *Tamiasciurus hudsonicus* | Terrestrial | 1 | 1 | (109) |
|  | *Thalassarche melanophris* | Terrestrial | 1 |  | (110, 111) |
|  | *Umbonium costatum* | Marine | 8 | 8 | (112) |
|  | *Urocitellus armatus* | Terrestrial | 1 | 1 | (113) |
|  | *Urocitellus beldingi* | Terrestrial | 1 |  | (114) |
|  | *Urocyon littoralis* | Terrestrial | 2 | 2 | (115) |
|  | *Ursus americanus* | Terrestrial | 4 | 4 | (116–118) |
|  | *Ursus arctos horribilis* | Terrestrial | 1 |  | (119) |
|  | *Ursus maritimus* | Terrestrial | 5 | 5 | (120) |
|  | *Vipera aspis* | Terrestrial | 1 | 1 | (121) |
|  | *Xenosaurus grandis* | Terrestrial | 4 | 4 | (122) |
|  | *Xenosaurus platyceps* | Terrestrial | 6 | 6 | (123) |
|  | *Xestospongia muta* | Marine | 5 | 2 | (124) |
|  | *Zalophus californianus* | Marine | 9 | 7 | (125) |
|  | *Zoarces viviparus* | Marine | 4 | 1 | (126) |
|  | *Zootoca vivipara* | Terrestrial | 1 |  | (127) |
|  | *Abies balsamea* | Terrestrial | 2 |  | (128) |
|  | *Abies concolor* | Terrestrial | 12 | 11 | (129) |
|  | *Abies homolepis* | Terrestrial | 1 |  | (130) |
|  | *Abies magnifica* | Terrestrial | 11 | 9 | (129) |
|  | *Abies sachalinensis* | Terrestrial | 3 | 2 | (131, 132) |
|  | *Acacia aneura* | Terrestrial | 1 |  |  |
|  | *Acacia victoriae* | Terrestrial | 1 |  | (133) |
|  | *Acer palmatum* | Terrestrial | 2 | 2 | (134) |
|  | *Acer pictum* | Terrestrial | 2 | 2 | (134) |
|  | *Acer rufinerve* | Terrestrial | 2 | 2 | (134) |
|  | *Acer saccharum* | Terrestrial | 6 | 6 | (135) |
|  | *Achnatherum calamagrostis* | Terrestrial | 3 | 3 | (136) |
|  | *Actaea cordifolia* | Terrestrial | 5 | 3 | (137) |
|  | *Actaea elata* | Terrestrial | 4 | 2 | (138, 139) |
|  | *Actaea spicata* | Terrestrial | 12 | 12 | (140) |
|  | *Adenocarpus gibbsianus* | Terrestrial | 13 | 2 | (141) |
|  | *Aesculus turbinata* | Terrestrial | 3 |  | (142) |
|  | *Agave marmorata* | Terrestrial | 2 |  | (143) |
|  | *Agave potatorum* | Terrestrial | 2 |  | (144) |
|  | *Agave vivipara* | Terrestrial | 2 |  | (145) |
|  | *Agrimonia eupatoria* | Terrestrial | 10 | 6 | (146) |
|  | *Agropyron cristatum* | Terrestrial | 2 |  | (147) |
|  | *Ailanthus altissima* | Terrestrial | 1 | 1 | (148, 149) |
|  | *Alliaria petiolata* | Terrestrial | 38 |  | (149–151) |
|  | *Allium monanthum* | Terrestrial | 2 | 1 | (152) |
|  | *Allium vineale* | Terrestrial | 2 |  | (149) |
|  | *Alnus rubra* | Terrestrial | 6 | 5 |  |
|  | *Ambrosia deltoidea* | Terrestrial | 1 | 1 |  |
|  | *Anarrhinum fruticosum* | Terrestrial | 5 | 3 | (141) |
|  | *Andira aubletii* | Terrestrial | 1 | 1 |  |
|  | *Andropogon gerardii* | Terrestrial | 20 | 9 | (153) |
|  | *Androsace vitaliana* | Terrestrial | 2 | 2 |  |
|  | *Anemone patens* | Terrestrial | 1 | 1 | (154) |
|  | *Anthericum liliago* | Terrestrial | 6 |  | (155) |
|  | *Anthericum ramosum* | Terrestrial | 11 |  | (155) |
|  | *Anthyllis vulneraria* | Terrestrial | 40 | 23 | (156, 157) |
|  | *Antirrhinum lopesianum* | Terrestrial | 3 | 2 | (141) |
|  | *Antirrhinum subbaeticum* | Terrestrial | 2 | 2 | (141) |
|  | *Aquilaria crassna* | Terrestrial | 1 | 1 | (158) |
|  | *Aquilegia chrysantha* | Terrestrial | 1 | 1 | (159) |
|  | *Aquilegia sp.* | Terrestrial | 1 |  | (160) |
|  | *Araucaria cunninghamii* | Terrestrial | 4 | 4 | (161) |
|  | *Araucaria muelleri* | Terrestrial | 4 | 4 | (162) |
|  | *Arctophila fulva* | Terrestrial | 3 |  | (163) |
|  | *Arenaria grandiflora* | Terrestrial | 5 | 3 | (141) |
|  | *Arisaema serratum* | Terrestrial | 1 | 1 | (164) |
|  | *Arisaema triphyllum* | Terrestrial | 4 |  | (165) |
|  | *Armeria maritima* | Terrestrial | 1 | 1 | (166) |
|  | *Armeria merinoi* | Terrestrial | 10 | 4 | (141) |
|  | *Arnica angustifolia* | Terrestrial | 1 |  | (167) |
|  | *Artemisia genipi* | Terrestrial | 4 | 4 | (157) |
|  | *Asclepias meadii* | Terrestrial | 2 | 2 | (168) |
|  | *Aspasia principissa* | Terrestrial | 1 | 1 | (169) |
|  | *Asplenium adulterinum* | Terrestrial | 18 |  | (170) |
|  | *Asplenium cuneifolium* | Terrestrial | 12 | 12 | (170) |
|  | *Asplenium scolopendrium* | Terrestrial | 6 | 6 | (171) |
|  | *Aster amellus* | Terrestrial | 27 |  | (172) |
|  | *Aster pyrenaeus* | Terrestrial | 5 |  | (141) |
|  | *Astragalus alopecurus* | Terrestrial | 15 | 6 |  |
|  | *Astragalus bibullatus* | Terrestrial | 4 | 4 | (173) |
|  | *Astragalus cremnophylax* | Terrestrial | 2 |  | (174) |
|  | *Astragalus michauxii* | Terrestrial | 1 | 1 | (175) |
|  | *Astragalus scaphoides* | Terrestrial | 115 | 101 | (176, 177) |
|  | *Astragalus tremolsianus* | Terrestrial | 5 | 3 | (141) |
|  | *Astragalus tyghensis* | Terrestrial | 45 | 36 | (138) |
|  | *Astrocaryum mexicanum* | Terrestrial | 7 | 5 | (178) |
|  | *Astrophytum asterias* | Terrestrial | 8 | 7 | (179) |
|  | *Astrophytum capricorne* | Terrestrial | 3 | 3 | (180) |
|  | *Astrophytum ornatum* | Terrestrial | 3 | 3 | (181) |
|  | *Atriplex acanthocarpa* | Terrestrial | 3 | 2 | (182) |
|  | *Atriplex canescens* | Terrestrial | 3 | 3 | (182) |
|  | *Aurinia saxatilis subsp. Saxatilis* | Terrestrial | 6 |  |  |
|  | *Avicennia germinans* | Terrestrial | 1 | 1 | (183) |
|  | *Balsamorhiza sagittata* | Terrestrial | 5 |  | (184) |
|  | *Banksia ericifolia* | Terrestrial | 1 | 1 | (185) |
|  | *Bertholletia excelsa* | Terrestrial | 2 | 2 | (186) |
|  | *Betula pubescens pumila* | Terrestrial | 3 |  | (187) |
|  | *Boechera fecunda* | Terrestrial | 14 | 13 | (188) |
|  | *Borassus aethiopum* | Terrestrial | 4 | 4 | (189) |
|  | *Bothriochloa insculpta* | Terrestrial | 4 |  | (190) |
|  | *Bouteloua rigidiseta* | Terrestrial | 3 |  | (191) |
|  | *Brassica insularis* | Terrestrial | 36 | 31 | (192) |
|  | *Brosimum alicastrum* | Terrestrial | 1 |  | (193) |
|  | *Bursera glabrifolia* | Terrestrial | 3 | 1 | (194) |
|  | *Calamagrostis canescens* | Terrestrial | 1 |  | (195) |
|  | *Calamus nambariensis* | Terrestrial | 1 |  | (196) |
|  | *Calamus rhabdocladus* | Terrestrial | 1 |  | (196) |
|  | *Calocedrus decurrens* | Terrestrial | 6 | 6 | (129) |
|  | *Calochortus albus* | Terrestrial | 2 | 2 | (197) |
|  | *Calochortus lyallii* | Terrestrial | 44 | 44 | (198) |
|  | *Calochortus obispoensis* | Terrestrial | 2 | 1 | (197) |
|  | *Calochortus pulchellus* | Terrestrial | 2 | 1 | (197) |
|  | *Calochortus tiburonensis* | Terrestrial | 2 | 2 | (197) |
|  | *Camellia japonica* | Terrestrial | 2 | 1 | (199) |
|  | *Carduus nutans* | Terrestrial | 5 | 4 | (151, 200) |
|  | *Carex bigelowii* | Terrestrial | 1 |  | (201) |
|  | *Carex humilis* | Terrestrial | 8 |  | (202) |
|  | *Carlina vulgaris* | Terrestrial | 8 | 3 |  |
|  | *Carnegiea gigantea* | Terrestrial | 1 | 1 | (203) |
|  | *Castanea dentata* | Terrestrial | 24 | 23 | (204) |
|  | *Catopsis compacta* | Terrestrial | 3 | 3 | (205) |
|  | *Catopsis sessiliflora* | Terrestrial | 2 | 1 | (206) |
|  | *Cecropia obtusifolia* | Terrestrial | 3 | 3 | (207) |
|  | *Cedrela odorata* | Terrestrial | 15 |  | (208) |
|  | *Centaurea horrida* | Terrestrial | 3 | 3 | (209) |
|  | *Centaurea jacea* | Terrestrial | 4 |  | (210) |
|  | *Cephalocereus senilis* | Terrestrial | 1 |  |  |
|  | *Chamaecrista lineata* | terrestrial | 12 | 11 | (211) |
|  | *Chamaedorea radicalis* | Terrestrial | 8 | 7 | (212) |
|  | *Cheirolophus metlesicsii* | Terrestrial | 5 | 2 | (141) |
|  | *Cherleria obtusiloba* | Terrestrial | 1 | 1 | (213) |
|  | *Chlorocardium rodiei* | Terrestrial | 1 | 1 | (214) |
|  | *Cirsium acaule* | Terrestrial | 2 | 2 | (148, 215) |
|  | *Cirsium dissectum* | Terrestrial | 13 |  | (216) |
|  | *Cirsium palustre* | Terrestrial | 3 |  | (217) |
|  | *Cirsium pannonicum* | Terrestrial | 1 | 1 | (215) |
|  | *Cirsium perplexans* | Terrestrial | 2 | 2 | (218) |
|  | *Cirsium pitcheri* | Terrestrial | 142 | 87 | (219, 220) |
|  | *Cirsium vulgare* | Terrestrial | 3 |  | (221–223) |
|  | *Cleistesiopsis bifaria* | Terrestrial | 7 |  | (224) |
|  | *Cleistesiopsis divaricata* | Terrestrial | 5 |  | (224) |
|  | *Clidemia hirta* | Terrestrial | 6 | 4 | (225) |
|  | *Coccothrinax readii* | Terrestrial | 1 | 1 | (226) |
|  | *Coespeletia spicata* | Terrestrial | 1 | 1 | (227) |
|  | *Coespeletia timotensis* | Terrestrial | 1 | 1 | (227) |
|  | *Conradina glabra* | Terrestrial | 4 | 3 | (228) |
|  | *Coprinopsis cinerea* | Terrestrial | 2 | 2 | (141) |
|  | *Corallorhiza trifida* | Terrestrial | 5 | 4 | (141) |
|  | *Cornus florida* | Terrestrial | 1 | 1 |  |
|  | *Cucurbita pepo* | Terrestrial | 3 | 3 | (229) |
|  | *Cynoglossum officinale* | Terrestrial | 1 |  | (230) |
|  | *Cypripedium calceolus* | Terrestrial | 30 |  | (231) |
|  | *Cypripedium fasciculatum* | Terrestrial | 24 | 23 |  |
|  | *Cypripedium lentiginosum* | Terrestrial | 1 |  | (232) |
|  | *Cyrtandra dentata* | Terrestrial | 20 | 20 | (233) |
|  | *Cytisus scoparius* | Terrestrial | 20 | 18 | (234) |
|  | *Dactylorhiza lapponica* | Terrestrial | 4 | 2 | (235, 236) |
|  | *Daemonorops poilanei* | Terrestrial | 1 |  | (196) |
|  | *Danthonia sericea* | Terrestrial | 10 | 8 | (237) |
|  | *Daphne rodriguezii* | Terrestrial | 41 |  | (238) |
|  | *Dendropanax trifidus* | Terrestrial | 1 |  | (199) |
|  | *Dicentra canadensis* | Terrestrial | 9 |  | (239) |
|  | *Dicerandra frutescens* | Terrestrial | 59 | 40 | (240) |
|  | *Dicorynia guianensis* | Terrestrial | 1 | 1 | (241) |
|  | *Dicymbe altsonii* | Terrestrial | 1 | 1 |  |
|  | *Digitaria eriantha* | Terrestrial | 4 |  | (190) |
|  | *Dioon caputoi* | Terrestrial | 3 | 3 | (242) |
|  | *Dioon edule* | Terrestrial | 3 |  | (243) |
|  | *Dioon merolae* | Terrestrial | 8 | 3 | (244) |
|  | *Dioon sonorense* | Terrestrial | 1 | 1 | (245) |
|  | *Dioon spinulosum* | Terrestrial | 2 | 2 |  |
|  | *Dioscorea chouardii* | Terrestrial | 1 | 1 | (246) |
|  | *Dipsacus fullonum* | Terrestrial | 2 |  | (247) |
|  | *Disporum sessile* | Terrestrial | 2 | 1 | (152) |
|  | *Disporum smilacinum* | Terrestrial | 2 | 1 | (152) |
|  | *Dracocephalum austriacum* | Terrestrial | 36 | 28 | (248) |
|  | *Duguetia neglecta* | Terrestrial | 1 | 1 |  |
|  | *Echeveria longissima* | Terrestrial | 1 |  | (249) |
|  | *Echinacea angustifolia* | Terrestrial | 8 | 8 | (250, 251) |
|  | *Echinocactus platyacanthus* | Terrestrial | 12 | 12 | (252) |
|  | *Echinospartum ibericum* | Terrestrial | 5 | 1 | (141) |
|  | *Encephalartos cycadifolius* | Terrestrial | 1 | 1 | (253) |
|  | *Encephalartos villosus* | Terrestrial | 1 | 1 | (253) |
|  | *Entandrophragma cylindricum* | Terrestrial | 1 | 1 | (254) |
|  | *Eperua falcata* | Terrestrial | 1 | 1 | (255) |
|  | *Epilobium latifolium* | Terrestrial | 2 |  | (256) |
|  | *Epipactis atrorubens* | Terrestrial | 11 | 3 | (257) |
|  | *Eremophila forrestii* | Terrestrial | 9 | 9 | (258) |
|  | *Eremophila maitlandii* | Terrestrial | 12 | 10 | (258) |
|  | *Eriogonum longifolium* | Terrestrial | 16 | 16 | (259) |
|  | *Eritrichium caucasicum* | Terrestrial | 4 | 2 | (260) |
|  | *Erodium paularense* | Terrestrial | 10 | 10 | (141) |
|  | *Erophila verna* | Terrestrial | 1 |  | (151) |
|  | *Erycina crista-galli* | Terrestrial | 2 | 2 |  |
|  | *Eryngium alpinum* | Terrestrial | 9 | 8 | (261) |
|  | *Eryngium cuneifolium* | Terrestrial | 48 | 38 | (262) |
|  | *Eryngium maritimum* | Terrestrial | 2 | 2 | (263) |
|  | *Erythranthe cardinalis* | Terrestrial | 12 | 10 | (264) |
|  | *Erythranthe lewisii* | Terrestrial | 12 | 8 | (264) |
|  | *Erythronium japonicum* | Terrestrial | 2 | 2 | (152, 265) |
|  | *Escobaria robbinsorum* | Terrestrial | 12 | 9 | (266) |
|  | *Escontria chiotilla* | Terrestrial | 2 | 2 |  |
|  | *Eupatorium perfoliatum* | Terrestrial | 3 |  | (267) |
|  | *Eupatorium resinosum* | Terrestrial | 3 |  | (267) |
|  | *Euphorbia fontqueriana* | Terrestrial | 5 | 4 | (141) |
|  | *Euterpe edulis* | Terrestrial | 3 | 3 | (268) |
|  | *Euterpe precatoria* | Terrestrial | 4 | 4 | (269) |
|  | *Fagus crenata* | Terrestrial | 1 |  | (130) |
|  | *Fagus grandifolia* | Terrestrial | 3 | 3 | (270) |
|  | *Festuca eskia* | Terrestrial | 2 |  | (271) |
|  | *Frasera speciosa* | Terrestrial | 34 |  | (272) |
|  | *Fritillaria biflora* | Terrestrial | 2 |  | (273) |
|  | *Fuscospora fusca* | terrestrial | 6 | 6 | (274) |
|  | *Gardenia actinocarpa* | Terrestrial | 4 | 2 | (275) |
|  | *Gentiana pneumonanthe* | Terrestrial | 4 | 2 | (276) |
|  | *Gentianella campestris* | Terrestrial | 1 |  | (277) |
|  | *Geonoma deversa* | Terrestrial | 4 |  | (278) |
|  | *Geonoma macrostachys* | Terrestrial | 1 | 1 | (279) |
|  | *Geonoma pohliana* | Terrestrial | 2 | 2 | (280) |
|  | *Geonoma schottiana* | Terrestrial | 4 | 2 | (281) |
|  | *Geranium sylvaticum* | Terrestrial | 3 | 3 | (282) |
|  | *Geum reptans* | Terrestrial | 4 |  | (283) |
|  | *Geum rivale* | Terrestrial | 6 | 3 | (146) |
|  | *Goeppertia ovandensis* | Terrestrial | 16 | 3 | (284) |
|  | *Grias peruviana* | Terrestrial | 1 | 1 | (193) |
|  | *Guaiacum sanctum* | Terrestrial | 4 | 2 |  |
|  | *Guarianthe aurantiaca* | Terrestrial | 3 | 3 | (285) |
|  | *Guettarda viburnoides* | Terrestrial | 3 | 3 | (286) |
|  | *Helenium virginicum* | Terrestrial | 1 | 1 | (287) |
|  | *Helianthemum juliae* | Terrestrial | 9 | 7 | (288) |
|  | *Helianthemum polygonoides* | Terrestrial | 5 | 5 | (141) |
|  | *Helianthemum teneriffae* | Terrestrial | 5 | 3 | (141) |
|  | *Helianthus divaricatus* | Terrestrial | 8 |  | (289) |
|  | *Heliconia acuminata* | Terrestrial | 9 | 9 | (290) |
|  | *Heteropogon contortus* | Terrestrial | 4 | 3 | (190) |
|  | *Heteropsis flexuosa* | Terrestrial | 2 |  | (291) |
|  | *Heteropsis macrophylla* | Terrestrial | 2 |  | (291) |
|  | *Heteropsis oblongifolia* | Terrestrial | 2 |  | (291) |
|  | *Hilaria mutica* | Terrestrial | 4 | 4 | (292) |
|  | *Himantoglossum hircinum* | Terrestrial | 1 | 1 | (293) |
|  | *Himatanthus drasticus* | Terrestrial | 8 | 8 | (294) |
|  | *Horkelia congesta* | Terrestrial | 6 |  |  |
|  | *Hudsonia montana* | Terrestrial | 1 |  | (295) |
|  | *Hydrastis canadensis* | Terrestrial | 29 |  | (296, 297) |
|  | *Hylocomium splendens* | Terrestrial | 12 |  | (298, 299) |
|  | *Hypericum cumulicola* | Terrestrial | 66 | 35 | (300) |
|  | *Hypochaeris radicata* | Terrestrial | 2 |  | (301) |
|  | *Iriartea deltoidea* | Terrestrial | 5 | 5 | (302) |
|  | *Isatis tinctoria* | Terrestrial | 1 |  | (303) |
|  | *Jacobaea vulgaris* | Terrestrial | 3 |  | (304) |
|  | *Jacquiniella leucomelana* | Terrestrial | 3 |  | (305) |
|  | *Jacquiniella teretifolia* | Terrestrial | 3 |  | (305) |
|  | *Juniperus procera* | Terrestrial | 1 | 1 | (306) |
|  | *Jurinea fontqueri* | Terrestrial | 5 | 3 | (141) |
|  | *Khaya senegalensis* | Terrestrial | 12 | 11 | (307) |
|  | *Knautia arvensis* | Terrestrial | 1 |  | (308) |
|  | *Kosteletzkya pentacarpos* | Terrestrial | 8 | 7 | (309) |
|  | *Lantana camara* | Terrestrial | 10 |  | (310) |
|  | *Laserpitium longiradium* | Terrestrial | 5 | 3 | (141) |
|  | *Lathyrus vernus* | Terrestrial | 32 | 17 | (311) |
|  | *Lechea cernua* | Terrestrial | 8 | 8 | (312) |
|  | *Lechea deckertii* | Terrestrial | 8 | 7 | (312) |
|  | *Leontodon saxatilis* | Terrestrial | 4 | 4 |  |
|  | *Lepidium davisii* | Terrestrial | 22 | 4 |  |
|  | *Leptocoryphium lanatum* | Terrestrial | 1 |  | (313) |
|  | *Lespedeza juncea sericea* | Terrestrial | 3 |  | (151, 314) |
|  | *Lespedeza virginica* | Terrestrial | 1 |  | (315) |
|  | *Liatris scariosa* | Terrestrial | 15 | 7 | (316) |
|  | *Ligularia sibirica* | Terrestrial | 33 | 6 | (317) |
|  | *Limonium carolinianum* | Terrestrial | 1 | 1 | (318) |
|  | *Limonium erectum* | Terrestrial | 5 | 5 | (141) |
|  | *Limonium malacitanum* | Terrestrial | 5 | 2 | (141) |
|  | *Linum catharticum* | Terrestrial | 1 |  | (319) |
|  | *Lomatium bradshawii* | Terrestrial | 9 | 9 | (138) |
|  | *Lomatium cookii* | Terrestrial | 10 | 9 | (138) |
|  | *Lonicera maackii* | Terrestrial | 2 | 2 | (149) |
|  | *Lophophora diffusa* | Terrestrial | 2 | 2 |  |
|  | *Lotus arinagensis* | Terrestrial | 6 |  | (141) |
|  | *Lupinus tidestromii* | Terrestrial | 9 | 9 | (320) |
|  | *Lycaste aromatica* | Terrestrial | 3 |  | (305) |
|  | *Machaerium cuspidatum* | Terrestrial | 3 |  | (321) |
|  | *Magnolia macrophylla* | Terrestrial | 3 | 3 | (322) |
|  | *Mammillaria crucigera* | Terrestrial | 2 | 2 | (323) |
|  | *Mammillaria dixanthocentron* | Terrestrial | 1 | 1 |  |
|  | *Mammillaria gaumeri* | Terrestrial | 24 |  |  |
|  | *Mammillaria hernandezii* | Terrestrial | 6 | 5 |  |
|  | *Mammillaria huitzilopochtli* | Terrestrial | 5 | 5 | (324) |
|  | *Mammillaria magnimamma* | Terrestrial | 4 | 4 | (325) |
|  | *Mammillaria napina* | Terrestrial | 3 | 2 |  |
|  | *Mammillaria pectinifera* | Terrestrial | 1 | 1 | (326) |
|  | *Mammillaria solisioides* | Terrestrial | 3 | 3 |  |
|  | *Mammillaria supertexta* | Terrestrial | 1 | 1 |  |
|  | *Manilkara zapota* | Terrestrial | 2 | 2 | (327) |
|  | *Mauritia flexuosa* | Terrestrial | 1 | 1 | (328) |
|  | *Melaleuca viridiflora* | Terrestrial | 3 |  | (329) |
|  | *Melocactus bahiensis* | Terrestrial | 2 | 2 | (330) |
|  | *Melocactus ernestii* | Terrestrial | 4 | 4 |  |
|  | *Miconia albicans* | Terrestrial | 1 | 1 | (331) |
|  | *Miconia prasina* | Terrestrial | 8 |  | (332) |
|  | *Microberlinia bisulcata* | Terrestrial | 1 | 1 | (333) |
|  | *Mimulus guttatus* | Terrestrial | 22 |  | (334, 335) |
|  | *Molinia caerulea* | Terrestrial | 12 | 6 | (336) |
|  | *Myrsine guianensis* | Terrestrial | 1 |  | (331) |
|  | *Nardostachys jatamansi* | Terrestrial | 1 | 1 | (337) |
|  | *Neobuxbaumia macrocephala* | Terrestrial | 7 | 7 | (338, 339) |
|  | *Neobuxbaumia mezcalaensis* | Terrestrial | 6 | 6 | (338) |
|  | *Neobuxbaumia polylopha* | Terrestrial | 2 | 2 | (340) |
|  | *Neobuxbaumia tetetzo* | Terrestrial | 6 | 6 | (338, 339, 341) |
|  | *Neotinea ustulata* | Terrestrial | 5 | 5 | (342) |
|  | *Oenothera deltoides* | Terrestrial | 16 | 16 | (343) |
|  | *Olearia flocktoniae* | Terrestrial | 8 |  | (344) |
|  | *Oncidium poikilostalix* | Terrestrial | 2 | 2 | (345, 346) |
|  | *Opuntia macrocentra* | Terrestrial | 2 |  | (347) |
|  | *Opuntia macrorhiza* | Terrestrial | 27 | 6 | (348) |
|  | *Opuntia microdasys* | Terrestrial | 4 |  |  |
|  | *Opuntia rastrera* | Terrestrial | 14 | 9 | (349) |
|  | *Orchis purpurea* | Terrestrial | 36 | 28 | (350) |
|  | *Oreocarya flava* | Terrestrial | 3 | 3 | (351) |
|  | *Oxalis acetosella* | Terrestrial | 6 |  | (352) |
|  | *Oxandra asbeckii* | Terrestrial | 1 | 1 | (255) |
|  | *Oxytropis jabalambrensis* | Terrestrial | 4 | 2 | (141) |
|  | *Pachycereus pecten-aboriginum* | Terrestrial | 3 | 3 | (353) |
|  | *Paeonia officinalis* | Terrestrial | 15 |  | (354) |
|  | *Paliurus ramosissimus* | Terrestrial | 2 |  | (355) |
|  | *Panax quinquefolius* | Terrestrial | 2 | 1 |  |
|  | *Parkinsonia aculeata* | Terrestrial | 1 | 1 | (356) |
|  | *Parolinia glabriuscula* | Terrestrial | 5 | 4 | (141) |
|  | *Paronychia pulvinata* | Terrestrial | 1 | 1 | (213) |
|  | *Pediomelum esculentum* | Terrestrial | 8 | 6 |  |
|  | *Pentaclethra macroloba* | Terrestrial | 1 | 1 |  |
|  | *Periandra mediterranea* | Terrestrial | 1 | 1 | (357) |
|  | *Persoonia bargoensis* | Terrestrial | 4 | 3 |  |
|  | *Persoonia glaucescens* | Terrestrial | 3 | 3 |  |
|  | *Petrophile pulchella* | Terrestrial | 1 | 1 | (185) |
|  | *Phyllanthus emblica* | Terrestrial | 8 | 6 | (316, 358) |
|  | *Phyllanthus indofischeri* | Terrestrial | 3 | 3 | (358) |
|  | *Picea glehnii* | Terrestrial | 1 |  | (132) |
|  | *Picea jezoensis* | Terrestrial | 1 |  | (132) |
|  | *Pilosella floribunda* | Terrestrial | 1 |  | (359) |
|  | *Pinguicula alpina* | Terrestrial | 1 | 1 | (360) |
|  | *Pinguicula villosa* | Terrestrial | 1 | 1 | (360) |
|  | *Pinguicula vulgaris* | Terrestrial | 1 |  | (360) |
|  | *Pinus jeffreyi* | Terrestrial | 1 |  | (129) |
|  | *Pinus lambertiana* | Terrestrial | 6 | 6 | (129) |
|  | *Pinus maximartinezii* | Terrestrial | 1 | 1 | (361) |
|  | *Pinus nigra* | Terrestrial | 3 | 1 | (362) |
|  | *Pinus ponderosa* | Terrestrial | 1 | 1 | (129) |
|  | *Pinus strobus* | Terrestrial | 9 | 9 | (363) |
|  | *Pityopsis aspera* | Terrestrial | 2 | 1 | (364) |
|  | *Plantago coronopus* | Terrestrial | 35 | 29 | (365) |
|  | *Plantago lanceolata* | Terrestrial | 3 |  |  |
|  | *Platanthera hookeri* | Terrestrial | 4 |  | (366) |
|  | *Poa alpina* | Terrestrial | 6 |  | (157) |
|  | *Polemonium van-bruntiae* | Terrestrial | 9 |  | (367) |
|  | *Polygonum basiramium* | Terrestrial | 8 | 2 | (312) |
|  | *Potentilla anserina* | Terrestrial | 3 |  | (368) |
|  | *Potentilla recta* | Terrestrial | 1 |  | (369) |
|  | *Primula elatior* | Terrestrial | 21 | 13 | (370) |
|  | *Primula farinosa* | Terrestrial | 16 | 7 | (371) |
|  | *Primula veris* | Terrestrial | 4 | 1 | (372, 373) |
|  | *Primula vulgaris* | Terrestrial | 44 | 37 | (374) |
|  | *Prioria copaifera* | Terrestrial | 2 | 2 | (375) |
|  | *Prosartes lanuginosa* | Terrestrial | 4 | 4 | (376) |
|  | *Prosopis glandulosa* | Terrestrial | 4 | 4 | (377) |
|  | *Prosopis laevigata* | Terrestrial | 2 | 2 | (378) |
|  | *Prunus africana* | Terrestrial | 2 | 2 | (379) |
|  | *Prunus serotina* | Terrestrial | 3 | 2 | (380) |
|  | *Pseudomitrocereus fulviceps* | Terrestrial | 1 | 1 | (326) |
|  | *Pseudophoenix sargentii* | Terrestrial | 7 | 7 | (381) |
|  | *Pterocarpus angolensis* | Terrestrial | 1 | 1 |  |
|  | *Pterocereus gaumeri* | Terrestrial | 4 | 4 | (382) |
|  | *Ptychosperma macarthurii* | Terrestrial | 1 | 1 | (383) |
|  | *Purshia subintegra* | Terrestrial | 14 | 8 | (384) |
|  | *Pyrrocoma radiata* | Terrestrial | 85 | 65 | (138) |
|  | *Quercus mongolica* | Terrestrial | 1 | 1 | (131) |
|  | *Quercus rugosa* | Terrestrial | 1 | 1 |  |
|  | *Ramonda myconi* | Terrestrial | 15 | 13 | (385) |
|  | *Ranunculus acris* | Terrestrial | 2 |  | (386) |
|  | *Ranunculus peltatus* | Terrestrial | 5 | 3 |  |
|  | *Rhizophora mangle* | Terrestrial | 1 | 1 | (183) |
|  | *Rhododendron maximum* | Terrestrial | 3 | 1 | (387) |
|  | *Rhododendron ponticum* | Terrestrial | 20 | 4 | (388) |
|  | *Rhopalostylis sapida* | Terrestrial | 2 | 2 | (389) |
|  | *Rhus aromatica* | Terrestrial | 8 |  | (289) |
|  | *Rhus copallinum* | Terrestrial | 3 |  | (390) |
|  | *Rosa multiflora* | Terrestrial | 1 |  | (149) |
|  | *Rosmarinus tomentosus* | Terrestrial | 12 |  | (141) |
|  | *Roupala montana* | Terrestrial | 1 |  | (331) |
|  | *Rourea induta* | Terrestrial | 2 |  | (331) |
|  | *Rubus praecox* | Terrestrial | 3 |  | (391) |
|  | *Rubus saxatilis* | Terrestrial | 6 |  | (392) |
|  | *Rubus ursinus* | Terrestrial | 3 |  | (391) |
|  | *Rumex rupestris* | Terrestrial | 5 | 4 | (141) |
|  | *Ruppia maritima* | Terrestrial | 3 |  | (393) |
|  | *Sabal minor* | Terrestrial | 3 | 2 |  |
|  | *Salix arctica* | Terrestrial | 7 |  | (394) |
|  | *Santolina melidensis* | Terrestrial | 5 | 3 | (141) |
|  | *Saponaria bellidifolia* | Terrestrial | 14 | 5 | (395) |
|  | *Sarcocapnos baetica* | Terrestrial | 2 | 1 | (396) |
|  | *Sarcocapnos pulcherrima* | Terrestrial | 4 | 2 | (396) |
|  | *Sarracenia purpurea* | Terrestrial | 3 | 3 | (397, 398) |
|  | *Saussurea medusa* | Terrestrial | 4 |  | (399) |
|  | *Saxifraga aizoides* | Terrestrial | 4 | 3 | (157) |
|  | *Saxifraga cotyledon* | Terrestrial | 8 |  | (400) |
|  | *Scaphium macropodum* | Terrestrial | 6 | 6 | (401) |
|  | *Scorzonera hispanica* | Terrestrial | 1 | 1 | (402) |
|  | *Serapias cordigera* | Terrestrial | 39 | 24 | (403) |
|  | *Shorea leprosula* | Terrestrial | 3 | 3 | (404) |
|  | *Silene acaulis* | Terrestrial | 25 | 13 | (405) |
|  | *Silene ciliata* | Terrestrial | 7 | 5 | (406) |
|  | *Silene douglasii* | Terrestrial | 3 | 3 | (407) |
|  | *Silene spaldingii* | Terrestrial | 12 |  | (408) |
|  | *Solidago fistulosa* | Terrestrial | 3 | 1 |  |
|  | *Sonchus pustulatus* | Terrestrial | 1 | 1 | (409) |
|  | *Spartina alterniflora* | Terrestrial | 1 |  | (410) |
|  | *Spathoglottis plicata* | Terrestrial | 3 | 3 | (411) |
|  | *Stenocereus eruca* | Terrestrial | 15 | 3 |  |
|  | *Stryphnodendron microstachyum* | Terrestrial | 1 | 1 |  |
|  | *Succisa pratensis* | Terrestrial | 12 | 9 | (412, 413) |
|  | *Swallenia alexandrae* | Terrestrial | 1 | 1 | (414) |
|  | *Swietenia macrophylla* | Terrestrial | 1 | 1 | (415) |
|  | *Syngonanthus nitens* | Terrestrial | 15 |  | (416) |
|  | *Syzygium jambos* | Terrestrial | 1 | 1 | (417) |
|  | *Taraxacum campylodes* | Terrestrial | 1 |  | (149) |
|  | *Taraxacum erythrospermum* | Terrestrial | 2 |  | (149) |
|  | *Tetraberlinia bifoliolata* | Terrestrial | 1 | 1 | (333) |
|  | *Tetraneuris herbacea* | Terrestrial | 3 |  | (418) |
|  | *Thrinax radiata* | Terrestrial | 3 | 3 | (226) |
|  | *Thymus vulgaris* | Terrestrial | 4 | 1 | (141) |
|  | *Tillandsia brachycaulos* | Terrestrial | 3 |  | (419) |
|  | *Tillandsia deppeana* | Terrestrial | 2 | 2 | (305) |
|  | *Tillandsia juncea* | Terrestrial | 2 | 1 | (305) |
|  | *Tillandsia macdougallii* | Terrestrial | 5 | 5 | (420) |
|  | *Tillandsia multicaulis* | Terrestrial | 2 | 2 | (305, 421) |
|  | *Tillandsia punctulata* | Terrestrial | 2 | 2 | (305,421) |
|  | *Tillandsia violacea* | Terrestrial | 3 | 3 | (420) |
|  | *Tolumnia variegata* | Terrestrial | 1 |  | (422) |
|  | *Tragopogon pratensis* | Terrestrial | 1 |  | (423) |
|  | *Triadica sebifera* | terrestrial | 12 | 12 |  |
|  | *Trillium camschatcense* | Terrestrial | 1 | 1 | (424) |
|  | *Trillium grandiflorum* | Terrestrial | 46 | 41 | (425) |
|  | *Trillium ovatum* | Terrestrial | 23 | 2 |  |
|  | *Trillium persistens* | Terrestrial | 12 | 12 |  |
|  | *Trollius laxus* | Terrestrial | 11 | 10 | (426) |
|  | *Tsuga canadensis* | Terrestrial | 4 | 4 | (427) |
|  | *Vella pseudocytisus* | Terrestrial | 29 | 24 | (141) |
|  | *Verbascum fontqueri* | Terrestrial | 6 | 4 | (141) |
|  | *Verbascum thapsus* | Terrestrial | 1 |  | (151) |
|  | *Verticordia staminosa* | Terrestrial | 4 |  | (428) |
|  | *Vincetoxicum rossicum* | Terrestrial | 20 |  | (429) |
|  | *Viola elatior* | Terrestrial | 2 | 2 | (430) |
|  | *Viola pumila* | Terrestrial | 2 | 2 | (430) |
|  | *Viola sagittata* | Terrestrial | 1 | 1 | (431) |
|  | *Vriesea sanguinolenta* | Terrestrial | 4 | 4 | (432) |
|  | *Vulpicida pinastri* | Terrestrial | 6 | 6 | (433) |
|  | *Zamia amblyphyllidia* | Terrestrial | 2 |  | (434) |
|  | *Zamia inermis* | Terrestrial | 1 | 1 | (435) |
|  | *Zea diploperennis* | Terrestrial | 4 | 3 | (436) |
|  |  | **Total** | **3890** | **2242** |  |

**S2. Data cleaning**

**Table S2.** **The relative effect of data cleaning on our demographic and environmental variables.** Descriptive summary showing the number of outlying values omitted, the transformation format used to achieve normality, and the total number of populations missing estimates for each our demographic and environmental variables. Total number of populations is 2242 across all variables.

| **Variable** | **Omissions** | **Transformation** | **Missing*** |
| --- | --- | --- | --- |
| DAMPING RATIO (*ρ*) | 29 | *1/y^1.1^* | 29 |
| Survival | 112 | *NA* | 112 |
| Progression | 112 | *NA* | 112 |
| Retrogression | 114 | *NA* | 114 |
| Reproduction | 112 | *NA* | 112 |
| PERIOD OF OSCILLATION (*ψ*) | 33 | *1/y^0.4^* | 799 |
| Survival | 71 | *NA* | 837 |
| Progression | 71 | *NA* | 837 |
| Retrogression | 71 | *NA* | 837 |
| Reproduction | 68 | *NA* | 834 |
| REACTIVITY ($\bar{\rho}$) | 55 | *1/y^0.6^* | 55 |
| Survival | 112 | *log(\|y_max_\| – y)* | 113 |
| Progression | 60 | *log(\|y_max_\| – y)* | 60 |
| Retrogression | 69 | *log(y + \|y_min_\|)* | 69 |
| Reproduction | 81 | *log(\|y_max_\| – y)* | 81 |
| ATTENUATION ($\underline{\rho}$) | 0 | *y^0.7^* | 0 |
| Survival | 44 | *NA* | 45 |
| Progression | 60 | *log(\|y_max_\| – y)* | 60 |
| Retrogression | 41 | *NA* | 41 |
| Reproduction | 54 | *log(\|y_max_\| – y)* | 54 |
| MAXIMAL AMPLIFICATION ($\overline{\rho}_{max}$) | 55 | *1/y^0.5^* | 55 |
| Survival | 112 | *NA* | 113 |
| Progression | 68 | *log(\|y_max_\| – y)* | 68 |
| Retrogression | 112 | *NA* | 112 |
| Reproduction | 103 | *log(\|y_max_\| – y)* | 103 |
| MAXIMAL ATTENUATION ($\underline{\rho}_{max}$) | 0 | *y^0.4^* | 0 |
| Survival | 62 | *NA* | 63 |
| Progression | 87 | *log(\|y_max_\| – y)* | 87 |
| Retrogression | 63 | *log(y + \|y_min_\|)* | 63 |
| Reproduction | 89 | *NA* | 89 |
| FREQUENCY SPECTRUM  (Temperature, *β_T_*) | 7 | *NA* | 70 |
| AUTOCORRELATION  (Temperature, *a_T_*) | 37 | *NA* | 100 |
| THERMAL RANGE (*m*) | 1 | *NA* | 307 |
| FREQUENCY SPECTRUM  (Precipitation, *β_P_*) | 0 | *NA* | 63 |
| AUTOCORRELATION  (Precipitation, *a_P_*) | 0 | *NA* | 84 |
| **includes omitted values* | | | |

**S3. Constructing population-level phylogenetic trees**

A phylogenetic tree was constructed to ensure all our analyses accounted for covariance between closely related species. The scientific names of all unique species within our subset of Matrix Population Models (MPMs) extracted from the COMPADRE (1) and COMADRE databases (2), were cross-checked and taxonomically updated using the R package ‘*taxize*’ (437, 438). We used the R package ‘*rotl*’ (439), to extract phylogenetic data for each species from the Open tree of Life (OTL) (440). With this phylogenetic data, we constructed separate subtrees for brown/red algae, plants, and animal entities at the species level. We fused the three subtrees using the *bind.tree* tool from the ‘*phytools’* package (441). Whilst binding our subtrees, we combined the algae and plant subtrees first before then adding the animal subtree with marine sponges (*Demospongiae*) set as the outgroup.

We refined the structure of our species-level phylogenetic tree, specifically ensuring the tree was rooted and free of polytomies using the *is.rooted* and *multi2di* tools from the ‘*ape*’ package (442). Branch lengths were calculated using the Grafen method (443) assuming a Brownian motion mode of trait evolution, whereby the variance between species’ characteristics is directly proportional to time since divergence (444). The phylogenetic tree was then time-calibrated using the *chronos* function and checked to confirm ultrametricity, with any duplicated node labels renamed. Lastly, to accommodate intra-specific spatial variation in vital rates, we expanded this phylogenetic tree to include population-level information for the 257 species where data was available for more than one population (Fig. S1). For these repeated species, a number of artificial branches equal to the number of replicates, were bound to the corresponding species’ tip of the original phylogenetic tree. This process was carried out using the *bind.tip* function, with each of these artificial branches assigned an equal length of infinitesimally small value (*i.e*., 0.0000001). The branch lengths for our taxonomic tree were then used in all further analyses to ensure our findings accounted for ancestral relationships (445).

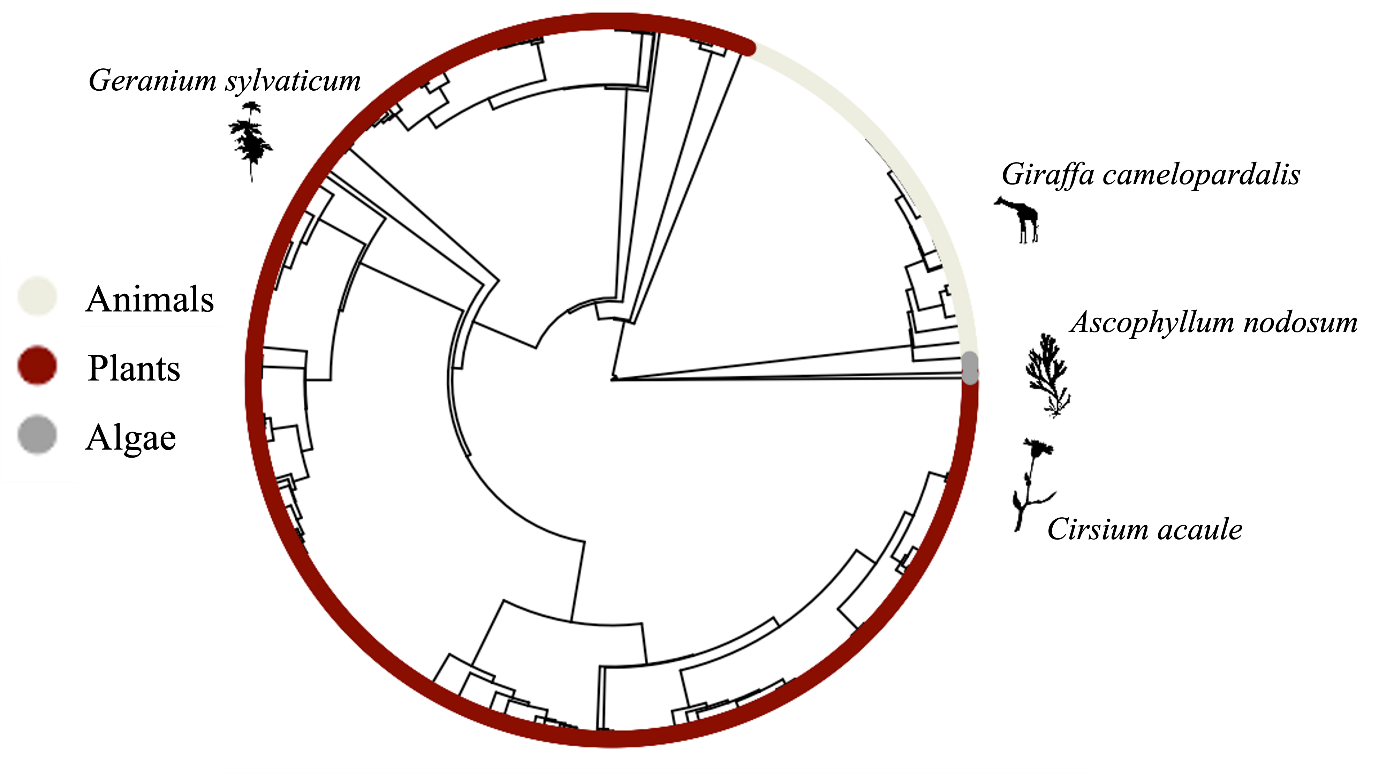

**Figure S1.** **Population-level phylogenetic tree displaying the relatedness of the 61 [37] animals, 305 [219] plants, and 3 [1] red/brown algae species used in this comparative assessment.** All our analyses have accounted for the phylogenetic signal between species. However, we have also allowed for the existence of multiple population entries for a number of species (shown above in square brackets), whilst assuming no within species trait variation.

**S4. Quantifying exposure to environmental stochasticity**

We quantified abiotic stochasticity to which each population has been exposed to examine the role of environmental stochasticity in shaping the resilience characteristics of *resistance*, *compensation*, and *recovery* across our 369 species. Using the GPS location information supplied with each extracted MPM from the COMPADRE (1) & COMADRE (2) databases, we linked each natural population to their corresponding abiotic environments.

To determine environmental stochasticity, we focused on maximum, and minimum monthly temperature (°C), and mean monthly precipitation records (kg m^-2^). These abiotic variables were selected as they are universally important across all ecoregions, except for in marine environments that are not directly affected by precipitation. Accordingly, we excluded MPMs associated with marine populations for this portion of our analysis (29 populations from 6 animal species, and 10 populations from 3 algal species). For our remaining 2184 terrestrial and 19 freshwater populations, we sourced high resolution (1 km^2^) monthly temperature and precipitation records from the CHELSA climate database (446). For each population, we extracted maximum, and minimum temperature readings, and mean precipitation records for a timeframe equal to the period during which the population was surveyed plus an additional 50 years prior to survey onset, to account for the effects of environmental legacy (447). Within our sample there was a total of 2 freshwater and 277 terrestrial populations for which no environmental data could be sourced. Subsequently, these 279 populations (12.7%) were excluded from our analyses into the role of environmental variance in shaping resilience attributes.

We used five metrics to quantify the extent of environmental variance imposed on each population: thermal autocorrelation (*a_T_*), thermal range (*m*), thermal frequency spectrum (*β_T_*), precipitation autocorrelation (*a_P_*), and precipitation frequency spectrum (*β_P_*). Extracted abiotic variables were arranged into time series depicting the 50+ year abiotic regimes to which each population was exposed. We then estimated the temporal autocorrelation of each temperature (*a_T_*) and precipitation (*a_P_*) time series, using the ‘*colorednoise*’ package (448). Next, we calculated the frequency spectrum of each time series. This metric is often referred to as the colour of environmental variation, and represented by a red to blue colour scale, with blue describing higher frequency variation and red variation dominated by low frequencies (449). The frequency spectrum of a time series is expressed by its spectral exponent (*β*), which is calculated as the negative slope coefficient of the linear regression between the log spectral density and log frequency of the time series (450). The spectral exponent of the temperature (*β_T_*) and precipitation (*β_P_*) regimes to which each population was exposed were calculated using the *spectrum* command from the ‘*stats*’ package (451). Finally, thermal range (*m*) was estimated as the mean difference between maximum and minimum monthly temperatures throughout a time series, providing a measure of the magnitude of any abiotic variation. Finally, prior to further analyses, outliers outside of the 95% confidence intervals were discarded for each of the aforementioned metrics of environmental stochasticity (Table S2), and each variable was checked for normality.

A Principal Components analysis (PCA) was used to explore the interrelationships between our five abiotic variables (Fig. S2), whilst we also evaluated their collinearity using variance-inflation factors (VIF). VIF reflects the degree to which, in a regression model, estimates of coefficients for any given variable are inflated by collinearity, with values of between 1 and 10 representing low collinearity (452, 453). VIF values were estimated using the *multicol* function from the ‘*fuzzySim*’ package (454). In our PCA the majority of the variation across our abiotic variables could be explained using the just the first two principal components (Proportional variance: PC1 0.43; PC2 0.28; PC3 0.16, PC4 0.08, PC5 0.05; Fig. S2). Here the first two principal components describe a gradient between the autocorrelation (*a_T_* & *a_P_*) and frequency spectrum (*β_T_* & *β_P_*) characteristics of abiotic environments (Table S3), reflecting a transition from red coloured environments characterised by positive autocorrelation (future abiotic conditions are conditional and similar to current conditions) and low frequency oscillations (seasonality), towards blue coloured environments with higher frequency oscillations, and negative temporal autocorrelation (i.e. future conditions contrast with current conditions). This trend corresponds with conditions expected of natural environments, as terrestrial environments are typically characterised by red coloured variation with marine environments considered even less variable (450, 455); hence the lack of blue coloured environments within our PCA (Fig. S2). However, with VIF confirming there was no collinearity among our five abiotic variables (VIF: *β_T_ =* 1.85; *a_T_* = 2.12; *m* = 1.17; *β_P_ =* 1.83; *a_P_* = 1.93), all variables were retained in further analyses.

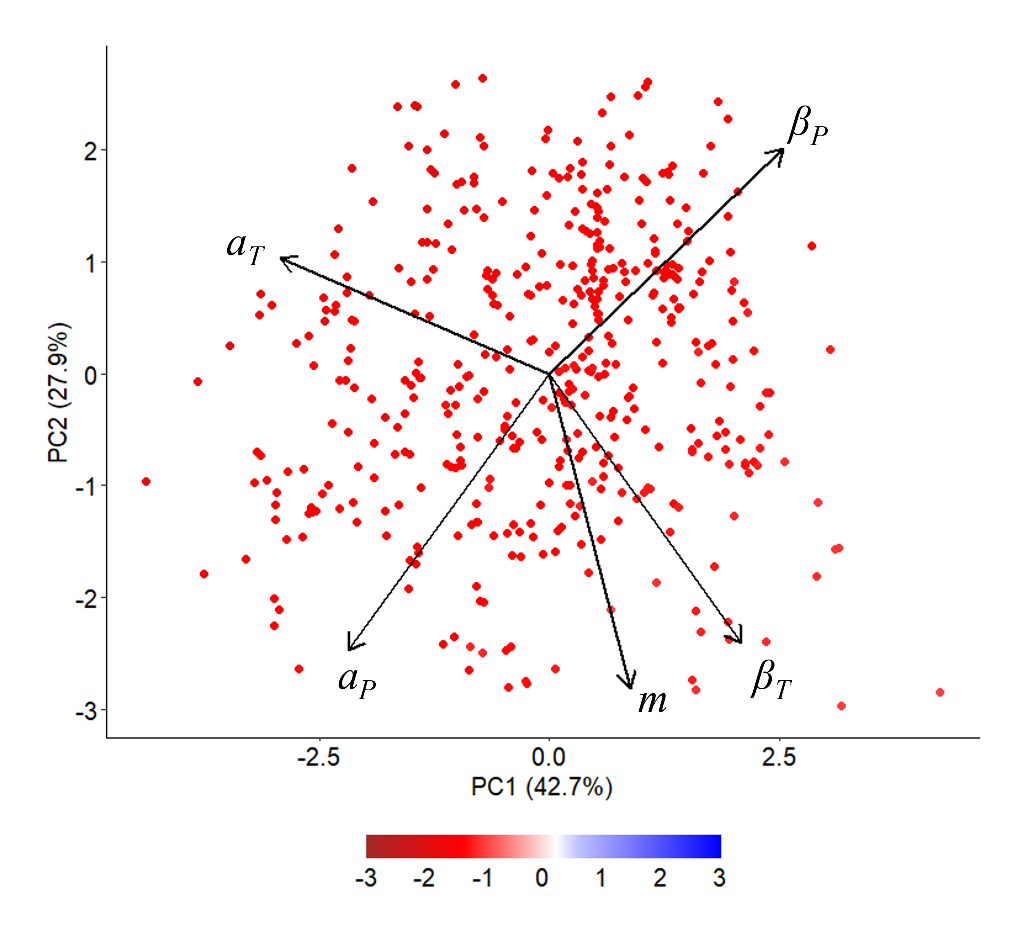

**Figure S2.** **Variation in the exposure of populations to environmental stochasticity corresponded with gradients in the autocorrelation and frequency spectrum characteristics of local abiotic regimes.** Principal component analysis (PCA) of the five metrics used to quantify exposure to environmental stochasticity: thermal autocorrelation (*a_T_*), thermal range (*m*), thermal frequency spectrum (*β_T_*), precipitation autocorrelation (*a_P_*), and precipitation frequency spectrum (*β_P_*) illustrating the degree of collinearity between the different variables. Colour scale depicts the gradient of environmental noise corresponding with transitions from red coloured environments characterised by positive autocorrelation and low frequency oscillations, towards blue coloured environments with higher frequency oscillations, and negative temporal autocorrelation. The colour of each environment was defined based on its associated thermal frequency exponent (*β_T_*) to demonstrate how abiotic variance regimes align with our five selected metrics.

**Table S3.** **Patterns within temperature and precipitation regimes characterised the relative exposure of populations to environmental stochasticity.** Principal component loadings of the five measures of environmental stochasticity, thermal autocorrelation (*a_T_*), thermal range (*m*), thermal frequency spectrum (*β_T_*), precipitation autocorrelation (*a_P_*), and precipitation frequency spectrum (*β_P_*) showing the relative influence of each abiotic variable across each principal component.

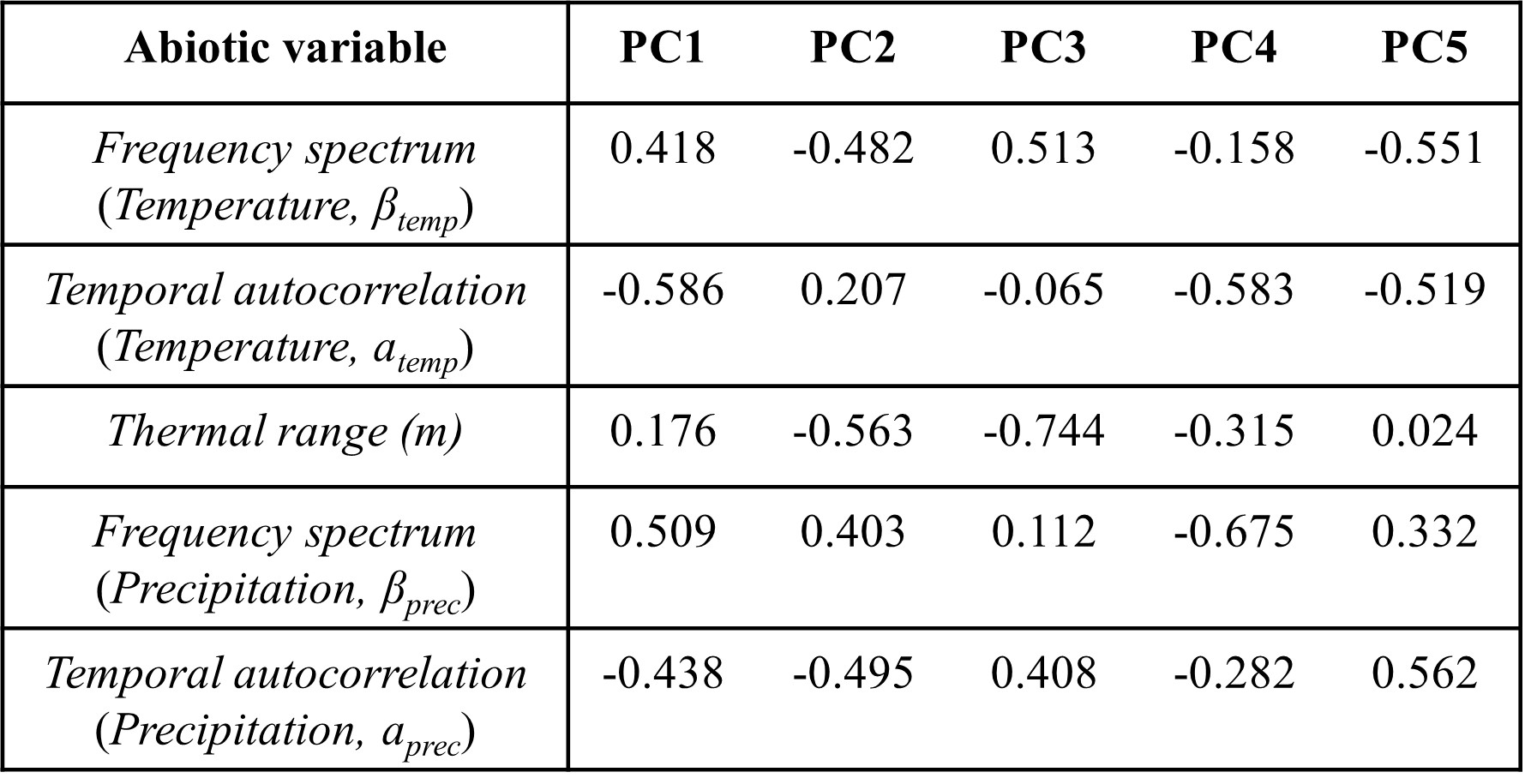

**S5. Phylogenetically imputed Partial Least Squares Regression analyses**

We initially carried out all pPLS analyses using only complete entries, omitting all populations missing estimates for any one variable, scaling and mean centring all predictor and response variables in each case. Across our dataset, no variable was missing from more than 6% of populations, except for the five variables describing the period of oscillation and its vital rate sensitivities, which were missing in 35-38% of populations (Table S2). To maximise our sample size (*n*) across each regression analysis, we omitted populations with incomplete entries separately across subsets of our data relating to each transient metric ($n_{\rho}$ = 1969; $n_{\psi}$ = 1263; $n_{\overline{\text{ρ}}}$ = 2055; $n_{\overline{\rho}max}$ = 2017; $n_{\underline{\text{ρ}}}$ = 2044; $n_{\underline{\rho}max}$ = 1988). However, we also repeated each analysis, with missing entries across the demographic variables estimated using phylogenetic imputation. To impute missing values, we first calculated the phylogenetic signal (Pagel’s *λ* [456]) of each transient and sensitivity variable using the *phylosig* function from the ‘*phytools*’ package (441). Pagel’s *λ* exists on the scale 0 < *λ* > 1, with 0 indicating traits have evolved independently of phylogeny, and 1 representing a high phylogenetic signal. Next, for all variables exhibiting a strong phylogenetic signal (Pagel’s *λ* ≥ 0.65) that differed significantly from 0 (*p* < 0.05), missing entries were imputed assuming a Brownian motion evolutionary model using the *phylopars* function of the ‘*Rphylopars*’ package (457).

Here we present the outputs of our regression analyses involving this imputed data as further evidence for any emerging patterns in the relationships between the transient dynamics of populations, their exposure to gradients in environmental stochasticity (Fig. S3), and their vital-rate sensitivities (Fig. S4). Our analysis using this imputed data displays congruent patterns to those reported using only complete entries. Indeed, our observations of limited association between gradients in environmental stochasticity and patterns in the transient dynamics of populations are maintained within the imputed data (Fig. S3; *r^2^ <* 0.006), whereas

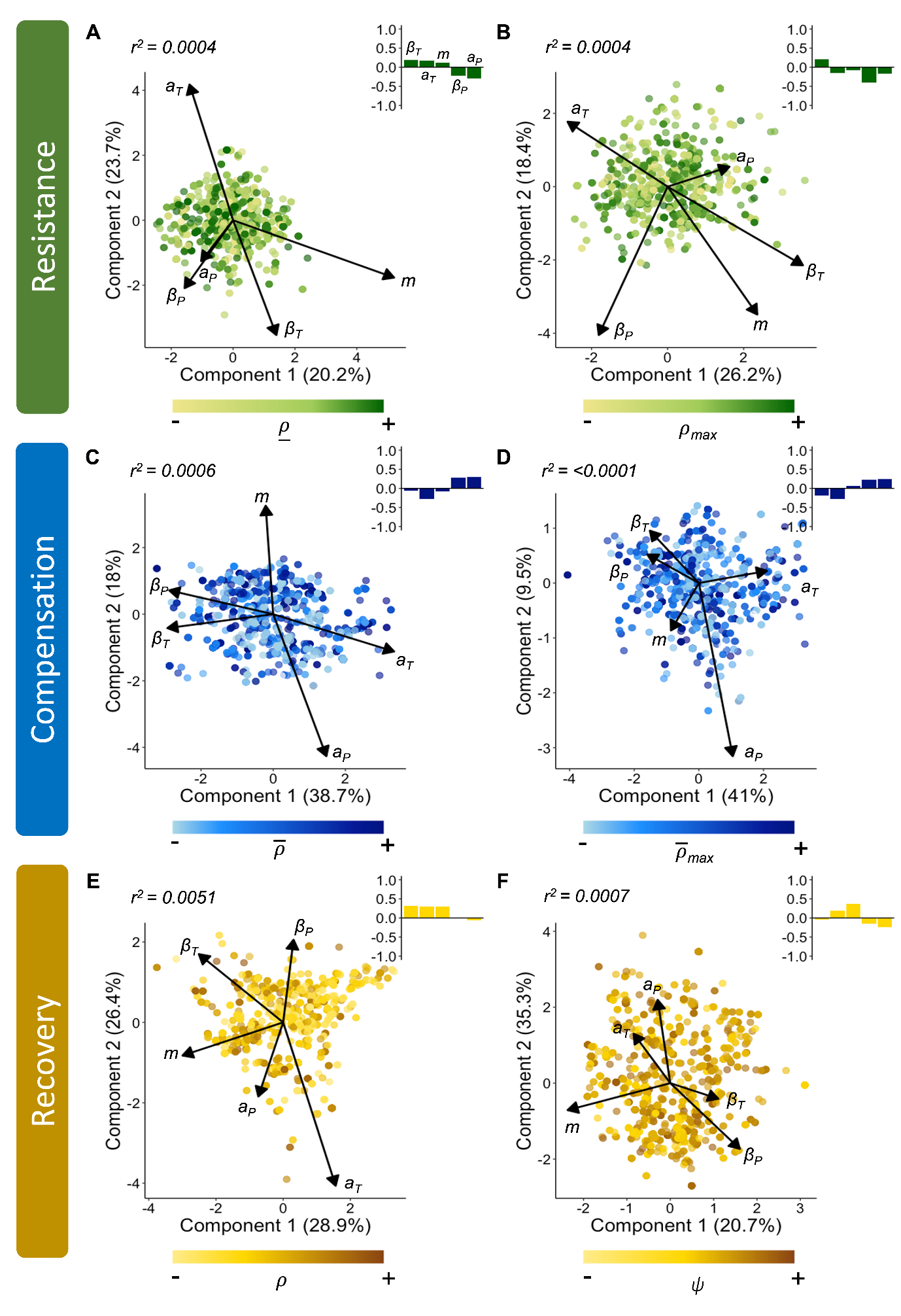

**Figure S3. The limited association between patterns across the demographic resilience attributes of resistance (green), compensation (blue), and recovery (orange) and the relative exposure of populations to environmental stochasticity is insensitive to phylogenetic imputation.** Scores and loadings from a phylogenetically-weighted Partial Least Squares regression analysis exploring the correlation between patterns in the variation of the six transient metrics of **(A)** first-step attenuation ($\underline{\rho}$), **(B)** maximal attenuation ($\underline{\rho}_{max}$), **(C)** reactivity ($\overline{\rho}$), **(D)** maximal amplification ($\overline{\rho}_{max}$), **(E)** damping ratio ($\rho$), and **(F)** period of oscillation (*ψ*), and our five measures of environmental stochasticity, temperature frequency spectrum (*β_T_*), temperature autocorrelation (*a_T_*), thermal range/magnitude (*m*), precipitation frequency spectrum (*β_P_*), and precipitation autocorrelation (*a_P_*) using a dataset with missing entries estimated through phylogenetic imputation. Colour gradation reflects the relative magnitude of each transient metric estimated for each population, with darker shades indicating higher estimates. Insert barplots are the standardised regression coefficients (*b*) highlighting the relative weighting of each abiotic variable in the overall capacity of each model to explain variation within each transient metric (*r^2^*).

the predictive capacity of our imputed vital rate sensitivity variables remains almost identical to those originally reported (*r^2^*; Fig S4). We note here that whilst each transient metric relating to the resilience attributes of resistance and compensation, and their vital rate sensitivities all displayed a strong phylogenetic signal (Pagel’s *λ* > 0.94, *p* <0.001; *see results*), this was not the case for our measures of demographic recovery. Both the transient metrics of damping ratio and period of oscillation displayed strong phylogenetic signal (Pagel’s *λ*: *ρ* = 0.996; *ψ* = 0.992; *p* <0.001 in both cases), but their vital rate sensitivities did not (*see results*). Subsequently, we were only able to examine patterns in the demographic selection pressures of recovery using complete entries.

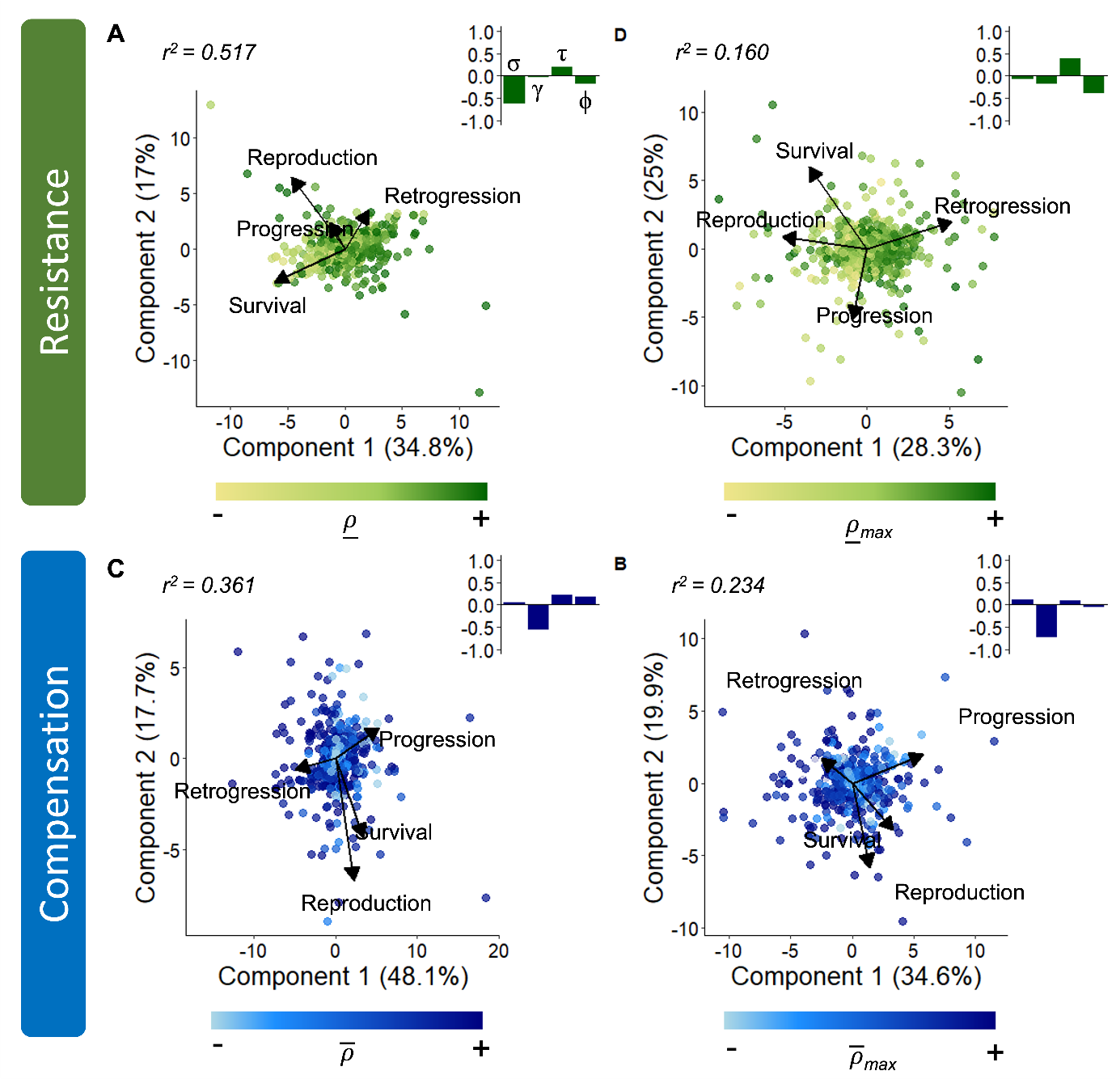

**Figure S4. Patterns within the vital rate sensitivities of the resilience attributes of resistance (green) and compensation (blue) are insensitive to phylogenetic imputation.** Scores and loadings from Partial Least Squares regression analysis of the sensitivity patterns of the four metrics of transient dynamics **(A)** first-step attenuation ($\underline{\rho}$), **(B)** maximal attenuation ($\underline{\rho}_{max}$), **(C)** reactivity ($\overline{\rho}$), and **(D)** maximal amplification ($\overline{\rho}_{max}$) with regards to the vital rates of survival (*σ*), progression (*γ*), retrogression (*τ*), and reproduction (*ϕ*) using a dataset with missing demographic entries phylogenetically imputed. Note that the transient metrics of damping ratio and period of oscillation have been excluded from this analysis due to a limited phylogenetic signal within the vital rate sensitivity variables of these two metrics. Colour gradation represents the magnitude of each transient metric recorded across each population, with darker shades indicating higher estimates. Insert barplots are the standardised regression coefficients (*b*) highlighting the relative weighting of each vital rate in the overall capacity of each model to explain variation within each transient metric (*r^2^*).

**S6. Accounting for population longevity**

We quantified the exposure of each population, within our sample, to environmental stochasticity using local temperature and precipitation records collected during the 50 years prior to the collection of any demographic data. However, the significance of any abiotic patterns experienced by each population during this 50-year window is likely contingent on their longevity. Across a 50-year period long-lived species, with generations spanning multiple decades, will experience fewer generations than shorter-lived species thereby diminishing the observable impact of any selection pressures on their trait characteristics (458). Thus, it was necessary we ensured that our capacity for exploring the selection pressures exerted by environmental stochasticity on the resilience attributes of natural populations was not inhibited by the inclusion of long-lived species.

To evaluate the influence of population longevity on our observations we repeated our phylogenetically weighted Partial Least Squares analyses evaluating the relationship between environmental stochasticity and the transient dynamics of populations using only short-lived species (Fig. S5). Each population was categorised as long- or short-lived according to its mean life expectancy (η_e_), which we estimated from its associated MPM using the R package ‘*IPMpack*’ (459). We then repeated our pPLS analyses using only populations for which η_e_ ≤ 10 years (n = 1606 populations). This threshold was selected to maximise the number of generations experienced by populations during our 50-year abiotic time series, whilst maintaining a suitable sample size for our analyses. Overall, whilst omitting longer-lived species did improve the predictive capacity our abiotic variables by an order of magnitude, the association between gradients in environmental variation and the resilience attributes of populations still remained negligible (*r^2^* < 0.001; Fig. S5). Indeed, the absolute magnitude of the Pearson’s coefficients (|*r|*) obtained when exploring the correlation between our measures of environmental stochasticity and transient dynamics all reflected a limited correlation
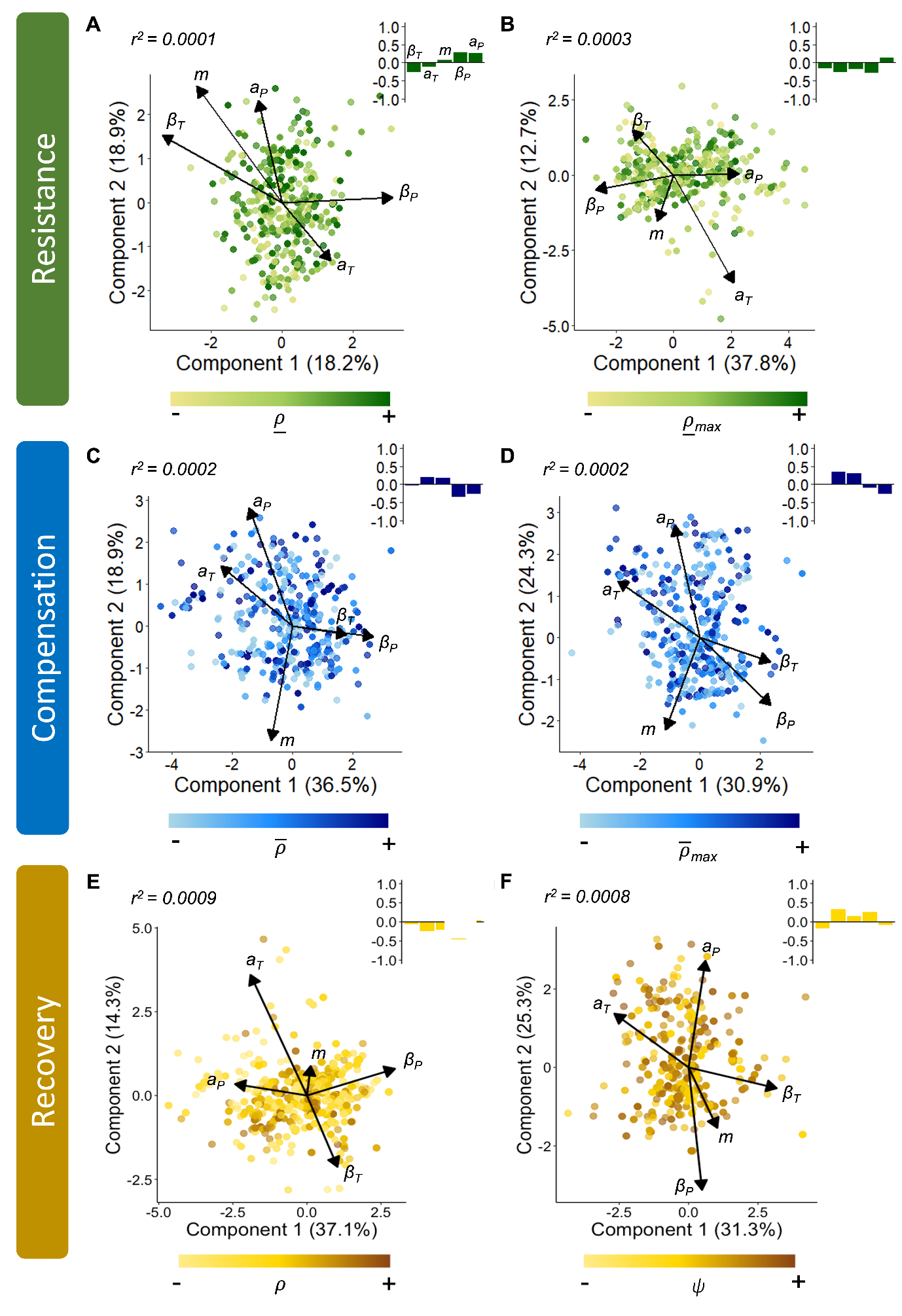
between our abiotic and demographic variables (*|r| <* 0.03; Table S4).

**Figure S5. The limited correlation between patterns across the demographic resilience attributes of resistance (green), compensation (blue), and recovery (orange) and the relative exposure of populations to environmental stochasticity is evident across short-lived populations.** Scores and loadings from a phylogenetically-weighted Partial Least Squares regression analysis exploring the correlation between patterns in the variation of the six transient metrics of **(A)** first-step attenuation ($\underline{\rho}$), **(B)** maximal attenuation ($\underline{\rho}_{max}$), **(C)** reactivity ($\overline{\rho}$), **(D)** maximal amplification ($\overline{\rho}_{max}$), **(E)** damping ratio ($\rho$), and **(F)** period of oscillation (*ψ*), and our five measures of environmental stochasticity: temperature frequency spectrum (*β_T_*), temperature autocorrelation (*a_T_*), thermal range/magnitude (*m*), precipitation frequency spectrum (*β_P_*), and precipitation autocorrelation (*a_P_*). Populations were selected for the analysis on the basis that they possess life expectancies of ≤ 10 years. Colour gradation reflects the relative magnitude of each transient metric estimated for each population, with darker shades indicating higher estimates. Insert barplots are the standardised regression coefficients (*b*) highlighting the relative weighting of each abiotic variable in the overall capacity of each model to explain variation within each transient metric (*r^2^*).

**Table S4. Patterns across the resilience attributes of resistance (green), compensation (blue), and recovery (orange) of short-lived populations do not correlate with their relative exposure to environmental stochasticity.** Using a phylogenetically-corrected Pearson’s test of correlation, we explored the correlation between the transient metrics of first-step attenuation ($\underline{\rho}$), maximal attenuation ($\underline{\rho}_{max}$), reactivity ($\overline{\rho}$), maximal amplification ($\overline{\rho}_{max}$), damping ratio ($\rho$), and period of oscillation (*ψ*), and each of our five measures of environmental stochasticity: temperature frequency spectrum (*β_T_*), temperature autocorrelation (*a_T_*), thermal range (*m*), precipitation frequency spectrum (*β_P_*), and precipitation autocorrelation (*a_P_*). Populations were selected for the analysis on the basis that they possessed life expectancies of ≤ 10 years.

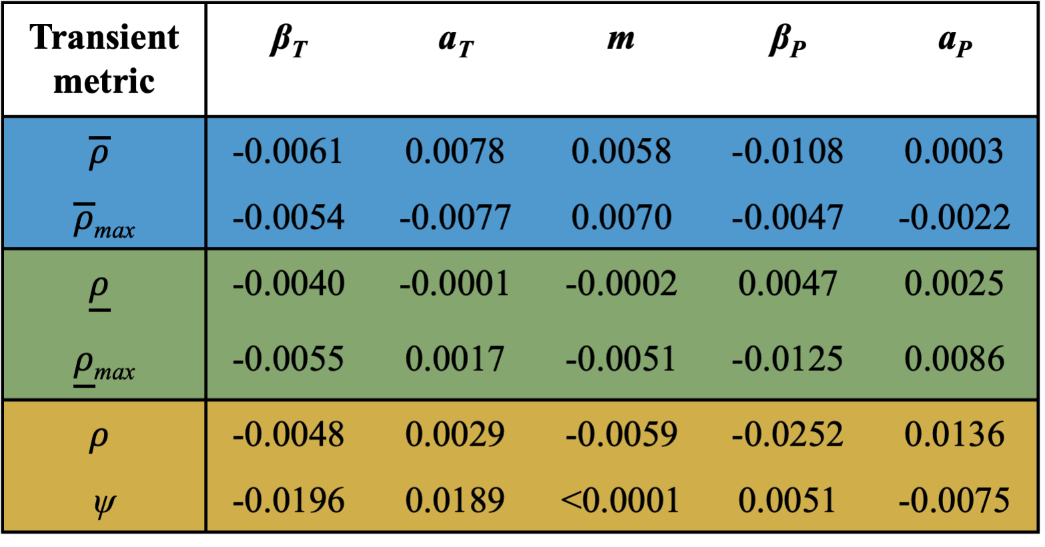

**S7. Accounting for legacy period length**

Evaluating how climate drivers influence demographic processes requires the identification of appropriate intervals of time during which climatic characteristics can be expected to impact upon vital rate properties (460). Accordingly, it was appropriate that we demonstrated that our observation, that the recent-past exposure of populations to environmental stochasticity does not predict their demographic resilience, is not sensitive to the breadth of time window (henceforth exposure legacy) used to quantify exposure to environmental stochasticity (Fig. S6 – Fig. S9). Originally, using the CHELSA climate database (446), we quantified exposure legacy across our population sample using high-resolution temperature and precipitation records covering the 50-years prior to demographic census (Supplementary S4). To illustrate the limited sensitivity of our observations to exposure legacy period length, we repeated our phylogenetically weighted Partial Least Squares analyses exploring the relationship between environmental stochasticity and the transient dynamics of populations using both 100-year (Fig. S6) and 5-year environmental legacy periods (Fig. S7). However, with the earliest CHELSA records dated from January 1901 (461), it was not possible for us to consistently estimate environmental legacy periods of 100-years across our entire population sample (which consisted of census start dates ranging from 1906 to 2016). Instead, to evaluate the relationship between environmental stochasticity and the transient dynamics of populations using 100-year environmental legacies we retained only studies for which it was possible to obtain 100-years of historical abiotic readings (915 populations). In both cases we demonstrate a neglible association between environmental stochasticity and the resilience attributes of populations (*r^2^* < 0.003; *|r| <* 0.04, Table S5 & Table S6). Moreover, this limited sensitivity continues to persist when considering only short-lived populations (*η_e_* ≤ 10; Supplementary S6) across both 100-year (Fig. S8 & Table S7) and 5-year environmental legacy periods (Fig. S9 & Table S8).

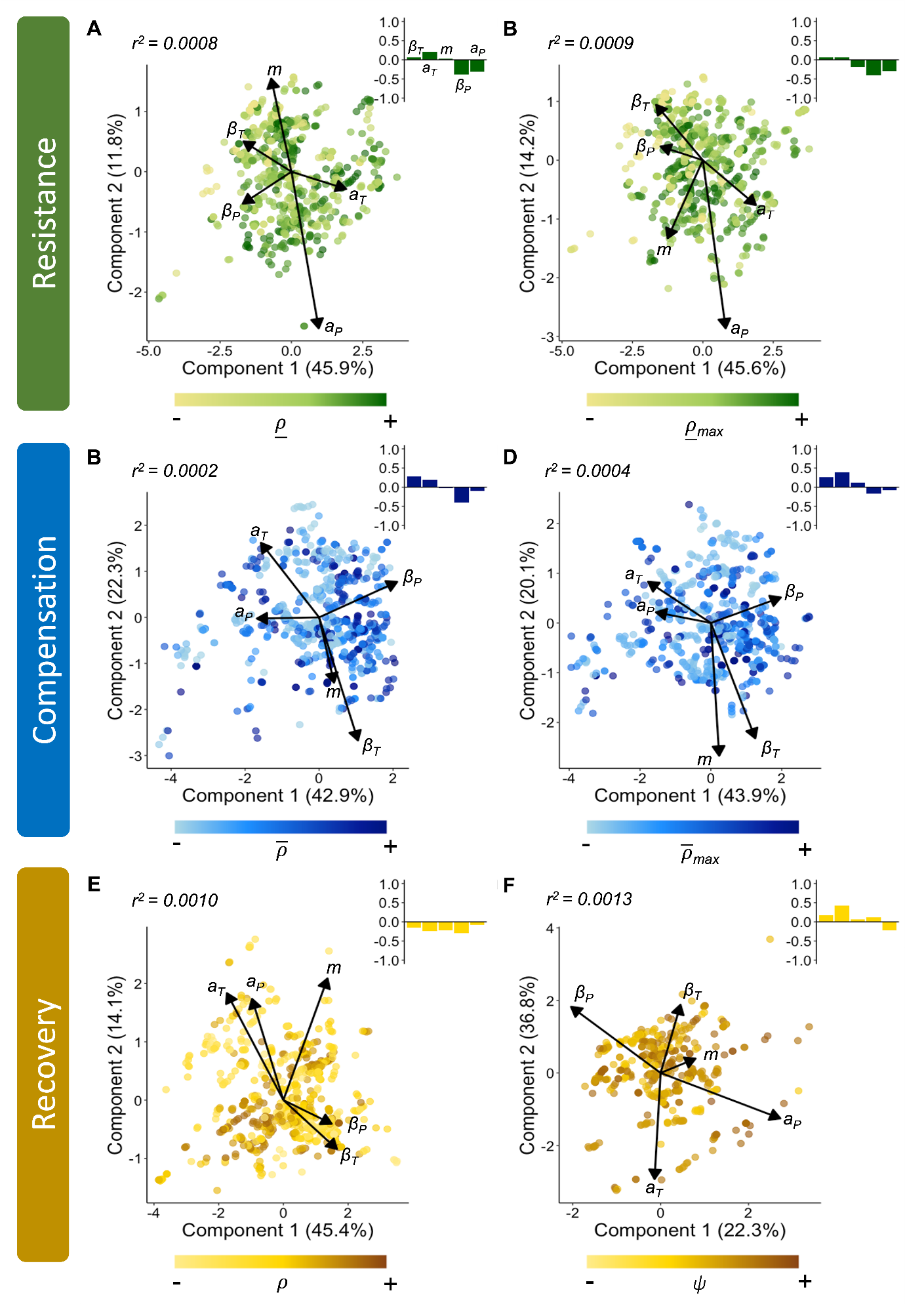

**Figure S6. The limited correlation between patterns across the demographic resilience attributes of resistance (green), compensation (blue), and recovery (orange) and the relative exposure of populations to environmental stochasticity persists when the exposure of populations is calculated using 100-year legacy periods.** Scores and loadings from a phylogenetically-weighted Partial Least Squares regression analysis exploring the correlation between patterns in the variation of the six transient metrics of **(A)** first-step attenuation ($\underline{\rho}$), **(B)** maximal attenuation ($\underline{\rho}_{max}$), **(C)** reactivity ($\overline{\rho}$), **(D)** maximal amplification ($\overline{\rho}_{max}$), **(E)** damping ratio ($\rho$), and **(F)** period of oscillation (*ψ*), and our five measures of environmental stochasticity: temperature frequency spectrum (*β_T_*), temperature autocorrelation (*a_T_*), thermal range/magnitude (*m*), precipitation frequency spectrum (*β_P_*), and precipitation autocorrelation (*a_P_*). Colour gradation reflects the relative magnitude of each transient metric estimated for each population, with darker shades indicating higher estimates. Insert barplots are the standardised regression coefficients (*b*) highlighting the relative weighting of each abiotic variable in the overall capacity of each model to explain variation within each transient metric (*r^2^*).

**Table S5. Patterns across the resilience attributes of resistance (green), compensation (blue), and recovery (orange) do not correlate with the exposure of populations to environmental stochasticity regimes over a 100-year period.** Using a phylogenetically-corrected Pearson’s test of correlation, we explored the correlation between the transient metrics of first-step attenuation ($\underline{\rho}$), maximal attenuation ($\underline{\rho}_{max}$), reactivity ($\overline{\rho}$), maximal amplification ($\overline{\rho}_{max}$), damping ratio ($\rho$), and period of oscillation (*ψ*), and each of our five measures of environmental stochasticity: temperature frequency spectrum (*β_T_*), temperature autocorrelation (*a_T_*), thermal range (*m*), precipitation frequency spectrum (*β_P_*), and precipitation autocorrelation (*a_P_*).

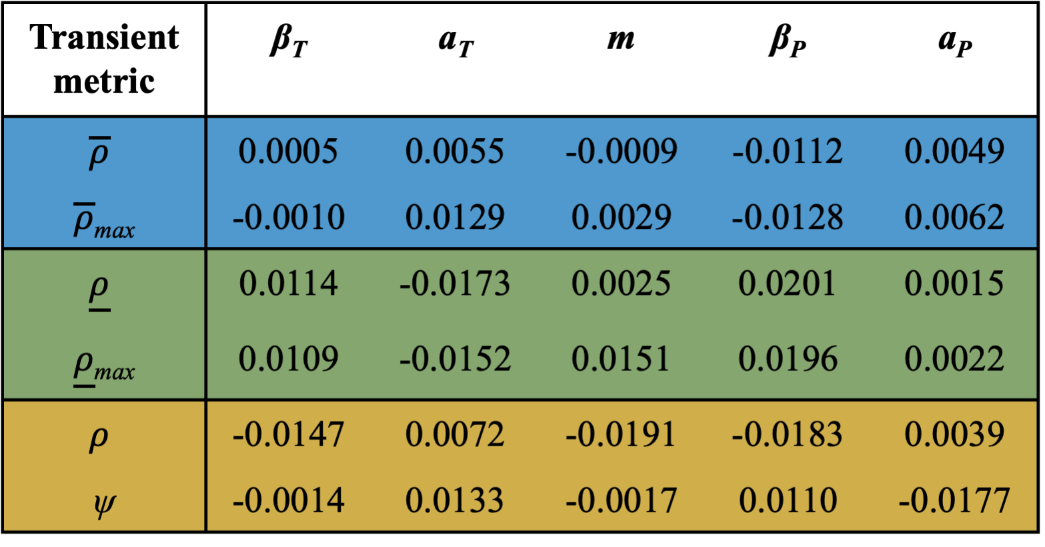

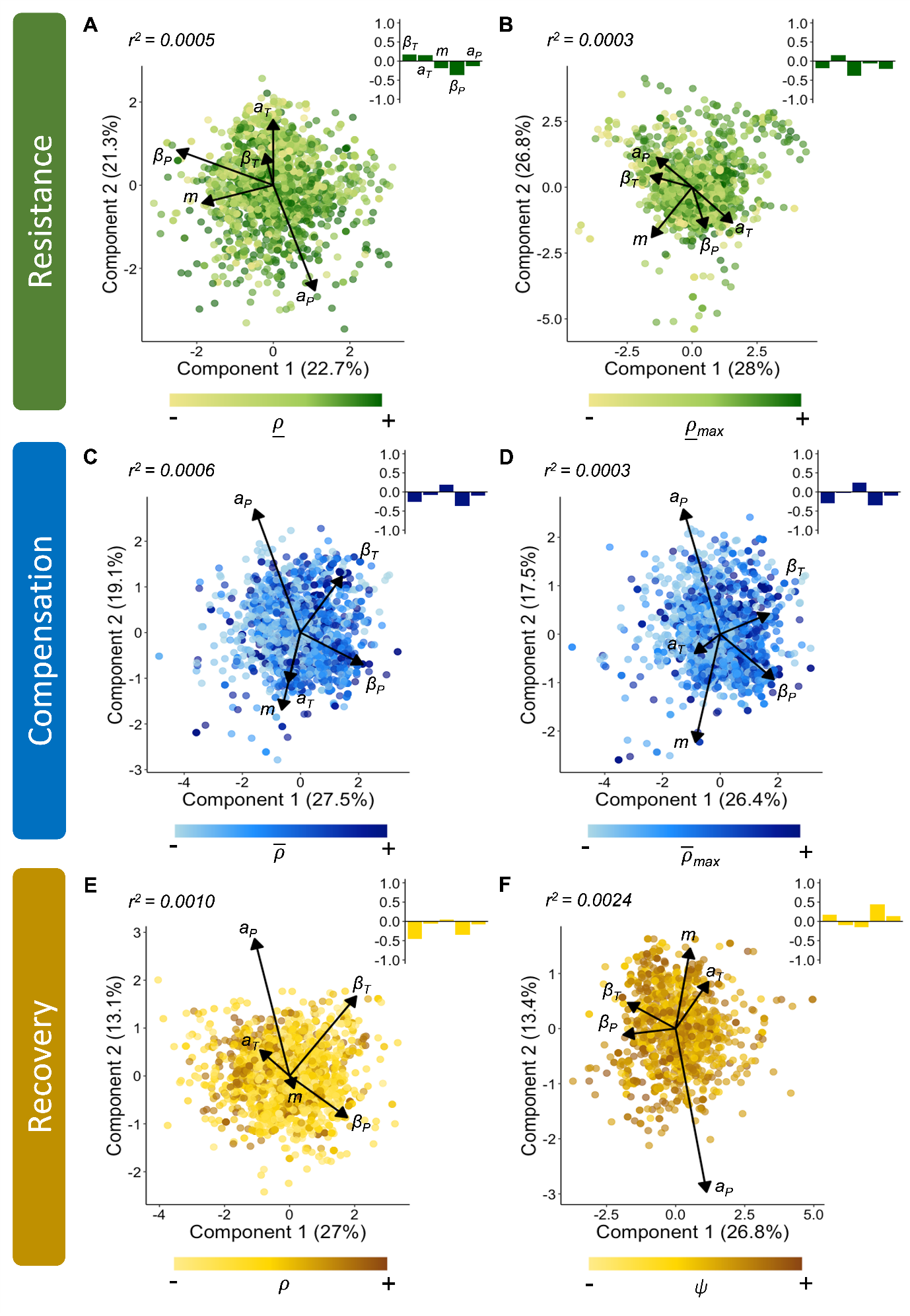

**Figure S7. The limited correlation between patterns across the demographic resilience attributes of resistance (green), compensation (blue), and recovery (orange) and the relative exposure of populations to environmental stochasticity persists when the exposure of populations is calculated using a 5-year legacy period.** Scores and loadings from a phylogenetically-weighted Partial Least Squares regression analysis exploring the correlation between patterns in the variation of the six transient metrics of **(A)** first-step attenuation ($\underline{\rho}$), **(B)** maximal attenuation ($\underline{\rho}_{max}$), **(C)** reactivity ($\overline{\rho}$), **(D)** maximal amplification ($\overline{\rho}_{max}$), **(E)** damping ratio ($\rho$), and **(F)** period of oscillation (*ψ*), and our five measures of environmental stochasticity: temperature frequency spectrum (*β_T_*), temperature autocorrelation (*a_T_*), thermal range/magnitude (*m*), precipitation frequency spectrum (*β_P_*), and precipitation autocorrelation (*a_P_*). Colour gradation reflects the relative magnitude of each transient metric estimated for each population, with darker shades indicating higher estimates. Insert barplots are the standardised regression coefficients (*b*) highlighting the relative weighting of each abiotic variable in the overall capacity of each model to explain variation within each transient metric (*r^2^*).

**Table S6. Patterns across the resilience attributes of resistance (green), compensation (blue), and recovery (orange) do not correlate with the exposure of populations to environmental stochasticity regimes over a 5-year period.** Using a phylogenetically-corrected Pearson’s test of correlation, we explored the correlation between the transient metrics of first-step attenuation ($\underline{\rho}$), maximal attenuation ($\underline{\rho}_{max}$), reactivity ($\overline{\rho}$), maximal amplification ($\overline{\rho}_{max}$), damping ratio ($\rho$), and period of oscillation (*ψ*), and each of our five measures of environmental stochasticity: temperature frequency spectrum (*β_T_*), temperature autocorrelation (*a_T_*), thermal range (*m*), precipitation frequency spectrum (*β_P_*), and precipitation autocorrelation (*a_P_*).

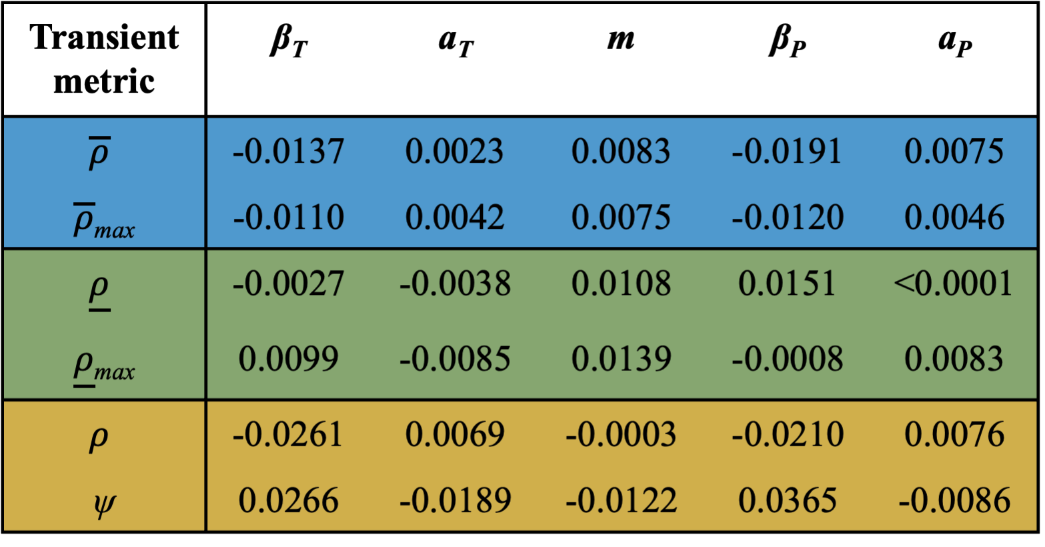

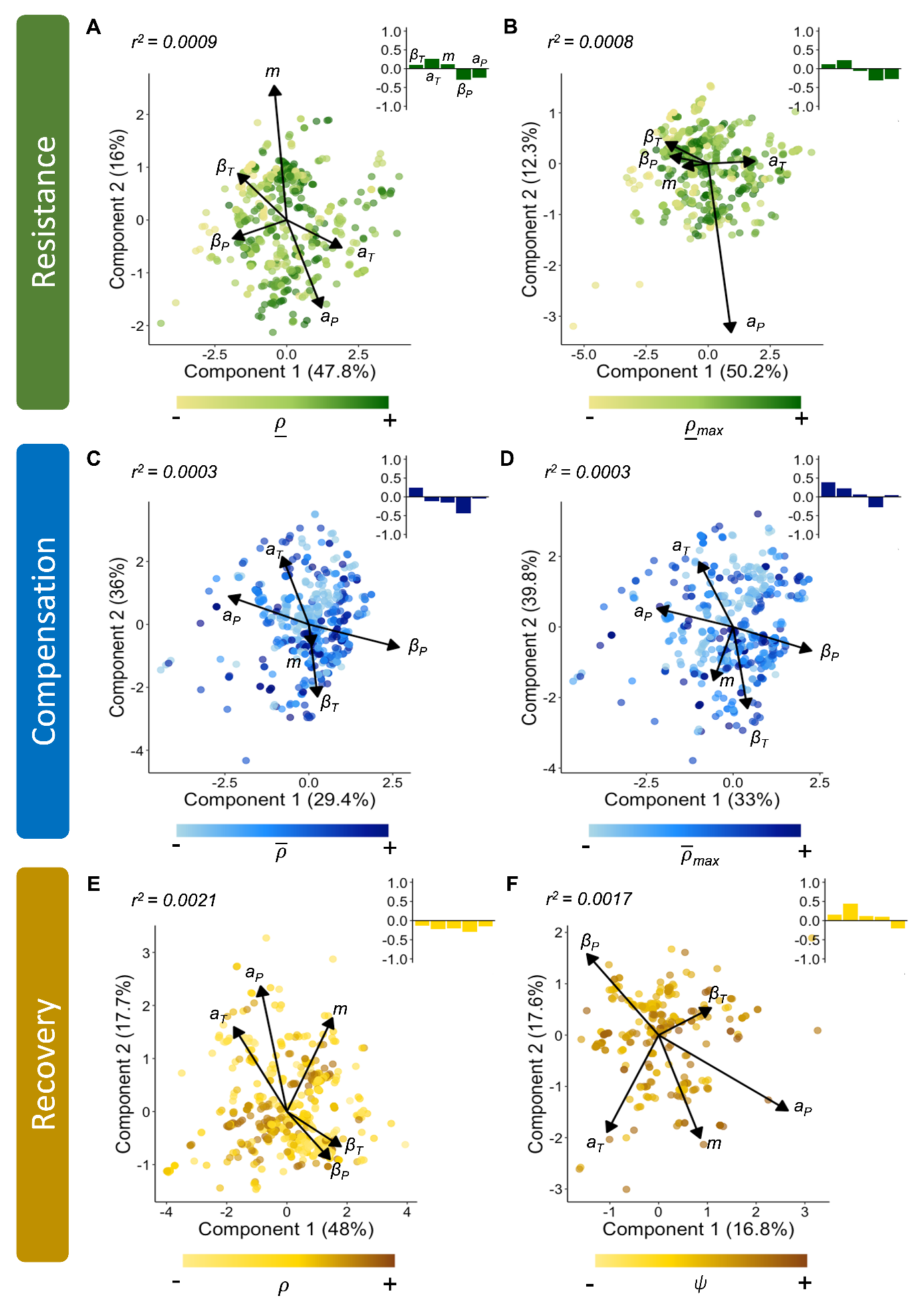

**Figure S8. The limited correlation between patterns across the demographic resilience attributes of resistance (green), compensation (blue), and recovery (orange) and the relative exposure of short-lived populations to environmental stochasticity remains when the exposure of populations is calculated using 100-year legacy periods.** Scores and loadings from a phylogenetically-weighted Partial Least Squares regression analysis exploring the correlation between patterns in the variation of the six transient metrics of **(A)** first-step attenuation ($\underline{\rho}$), **(B)** maximal attenuation ($\underline{\rho}_{max}$), **(C)** reactivity ($\overline{\rho}$), **(D)** maximal amplification ($\overline{\rho}_{max}$), **(E)** damping ratio ($\rho$), and **(F)** period of oscillation (*ψ*), and our five measures of environmental stochasticity: temperature frequency spectrum (*β_T_*), temperature autocorrelation (*a_T_*), thermal range/magnitude (*m*), precipitation frequency spectrum (*β_P_*), and precipitation autocorrelation (*a_P_*). Colour gradation reflects the relative magnitude of each transient metric estimated for each population, with darker shades indicating higher estimates. Insert barplots are the standardised regression coefficients (*b*) highlighting the relative weighting of each abiotic variable in the overall capacity of each model to explain variation within each transient metric (*r^2^*).

**Table S7. Patterns across the resilience attributes of resistance (green), compensation (blue), and recovery (orange) in short-lived populations do not correlate with their exposure to environmental stochasticity regimes over a 100-year period.** Using a phylogenetically-corrected Pearson’s test of correlation, we explored the correlation between the transient metrics of first-step attenuation ($\underline{\rho}$), maximal attenuation ($\underline{\rho}_{max}$), reactivity ($\overline{\rho}$), maximal amplification ($\overline{\rho}_{max}$), damping ratio ($\rho$), and period of oscillation (*ψ*), and each of our five measures of environmental stochasticity: temperature frequency spectrum (*β_T_*), temperature autocorrelation (*a_T_*), thermal range (*m*), precipitation frequency spectrum (*β_P_*), and precipitation autocorrelation (*a_P_*).

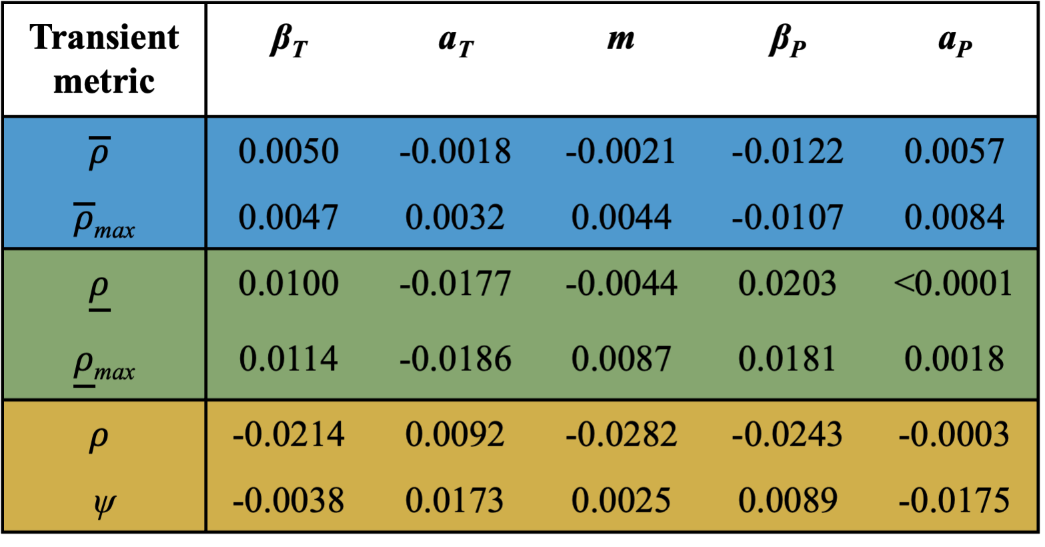

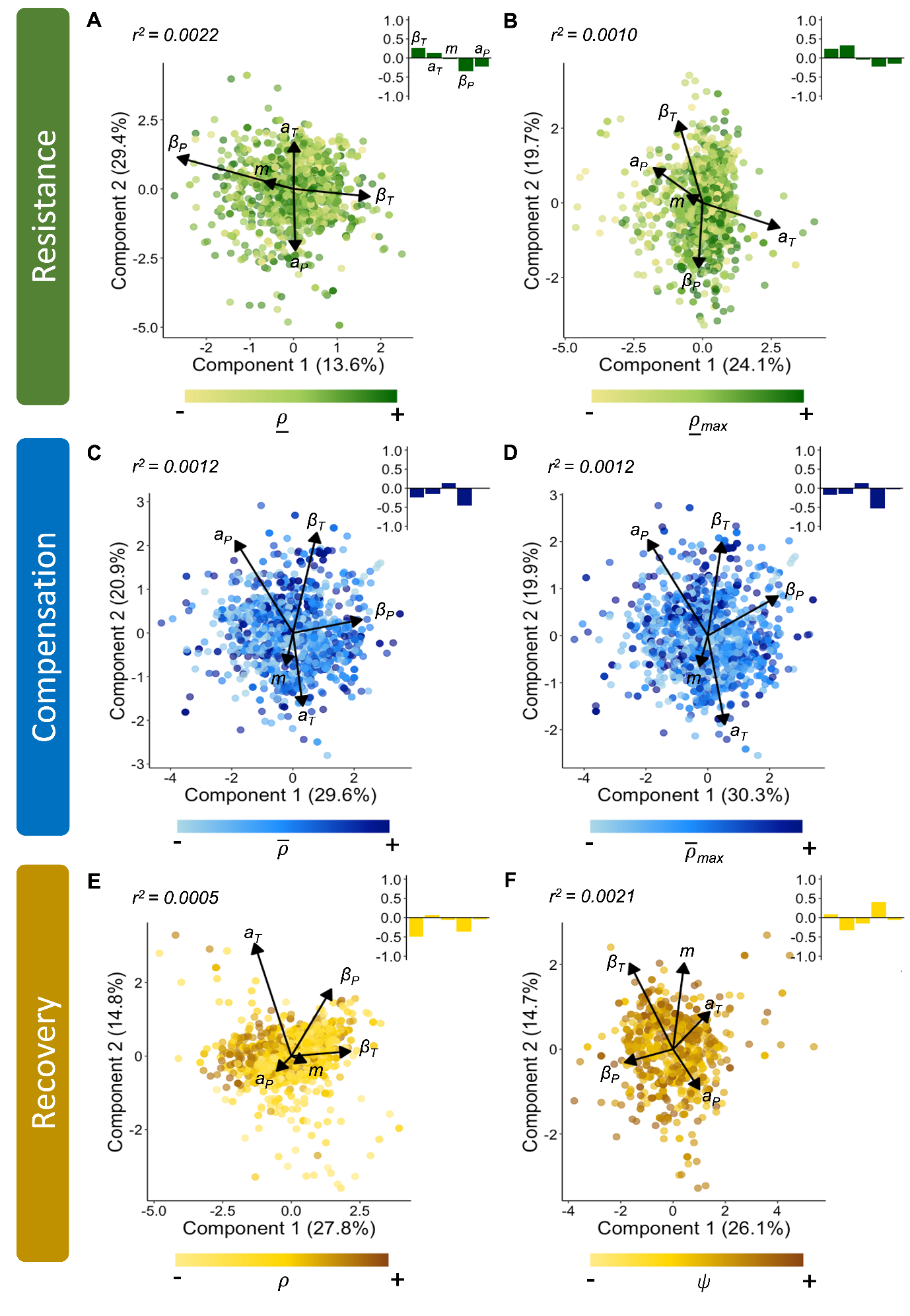

**Figure S9. The limited correlation between patterns across the demographic resilience attributes of resistance (green), compensation (blue), and recovery (orange) and the relative exposure of short-lived populations to environmental stochasticity remains when the exposure of populations is calculated using 5-year legacy periods.** Scores and loadings from a phylogenetically-weighted Partial Least Squares regression analysis exploring the correlation between patterns in the variation of the six transient metrics of **(A)** first-step attenuation ($\underline{\rho}$), **(B)** maximal attenuation ($\underline{\rho}_{max}$), **(C)** reactivity ($\overline{\rho}$), **(D)** maximal amplification ($\overline{\rho}_{max}$), **(E)** damping ratio ($\rho$), and **(F)** period of oscillation (*ψ*), and our five measures of environmental stochasticity: temperature frequency spectrum (*β_T_*), temperature autocorrelation (*a_T_*), thermal range/magnitude (*m*), precipitation frequency spectrum (*β_P_*), and precipitation autocorrelation (*a_P_*). Colour gradation reflects the relative magnitude of each transient metric estimated for each population, with darker shades indicating higher estimates. Insert barplots are the standardised regression coefficients (*b*) highlighting the relative weighting of each abiotic variable in the overall capacity of each model to explain variation within each transient metric (*r^2^*).

**Table S8. Patterns across the resilience attributes of resistance (green), compensation (blue), and recovery (orange) in short-lived populations do not correlate with their exposure to environmental stochasticity regimes over a 5-year period.** Using a phylogenetically-corrected Pearson’s test of correlation, we explored the correlation between the transient metrics of first-step attenuation ($\underline{\rho}$), maximal attenuation ($\underline{\rho}_{max}$), reactivity ($\overline{\rho}$), maximal amplification ($\overline{\rho}_{max}$), damping ratio ($\rho$), and period of oscillation (*ψ*), and each of our five measures of environmental stochasticity: temperature frequency spectrum (*β_T_*), temperature autocorrelation (*a_T_*), thermal range (*m*), precipitation frequency spectrum (*β_P_*), and precipitation autocorrelation (*a_P_*).

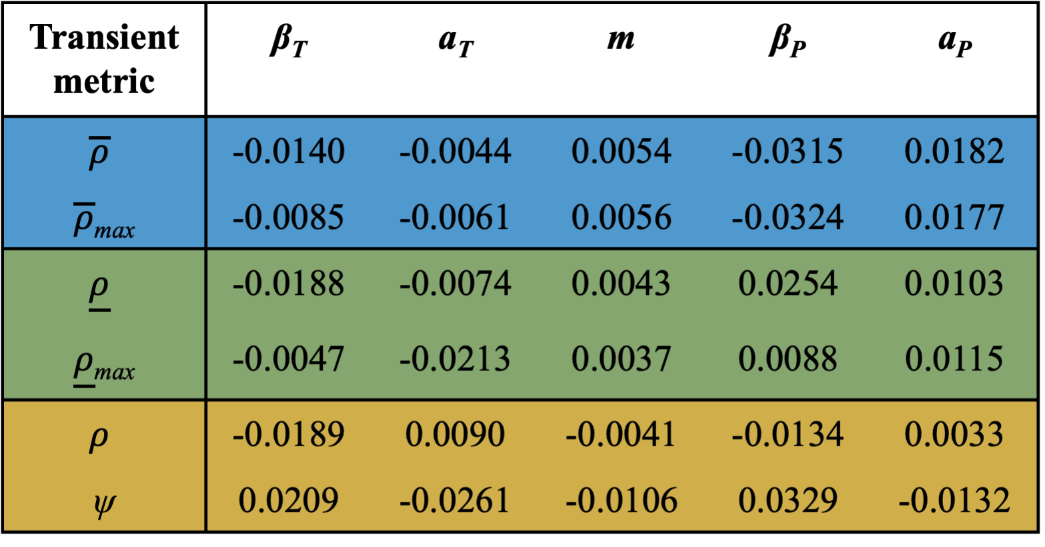
